# Increased aridity is associated with diversity and composition changes in the biocrust mycobiome

**DOI:** 10.1101/2025.03.05.641731

**Authors:** Kian Kelly, Xinzhan Liu, Jared Croyle, Jason E. Stajich

## Abstract

Drylands comprise 45% of Earth’s land area and contain ecologically critical soil surface communities known as biocrusts. Biocrusts are composed extremotolerant organisms including cyanobacteria, microfungi, algae, lichen, and bryophytes. Fungi in biocrusts help aggregate these communities and may form symbiotic relationships with nearby plants. Climate change threatens biocrusts, particularly moss biocrusts, but its effects on the biocrust mycobiome remain unknown. Here, we performed a culture-dependent and metabarcoding survey of the moss biocrust mycobiome across an aridity gradient to determine whether local climate influences fungal community composition. As the local aridity index increased, fungal communities exhibited greater homogeneity in beta diversity. At arid and hyper-arid sites, communities shifted toward more extremotolerant taxa. We identified a significant proportion of fungal reads and cultures from biocrusts that could not be classified. *Rhodotorula mucilaginosa* and *R. paludigena* were significantly enriched following surface sterilization of healthy biocrust mosses. This aligns with their known roles as plant endophytes. We also observed septate endophyte colonization in the photosynthetic tissues of mosses from arid climates. Collectively, these results suggest that the biocrust mycobiome will undergo significant shifts in diversity due to climate change, favoring extremotolerant taxa as climate conditions intensify. The survey results also highlight taxa with the potential to serve as bioinoculants for enhancing biocrust resilience to climate change. These findings offer valuable insights into the potential impacts of climate change on drylands and provide crucial information for biocrust conservation.

## Introduction

Drylands are the largest terrestrial biome, comprising 45% of earth’s land area and supporting 40% of the global population (Coleine et al., 2024). Because of their already extreme levels of aridity, lack of soil fertility and high UV radiation, drylands are particularly susceptible to climate change. Approximately one third of drylands are estimated to already be significantly degraded, which directly affects 250 million people and puts many others at risk (Coeline et al., 2024). California deserts are particularly vulnerable, as they are predicted to experience a temperature rise of 3% by the turn of the century (Cook et al., 2015). California is host to three major deserts: the Mojave Desert, the Sonoran Desert, and the Colorado Desert, each with variable levels of transpiration and precipitation.

Biological soil crusts (hereafter, biocrusts) are an example of cooperative communities within drylands (Pombubpa et al., 2019). Biocrusts are communities of bryophytes, lichen, cyanobacteria, and microfungi that aggregate the top milliliters of soil (Weber et al., 2022). Biocrusts cover approximately 12% of Earth’s terrestrial surface (Rodriguez-Caballero et al., 2018). Their net carbon uptake from the atmosphere, calculated as photosynthesis minus respiration, is estimated to be around 3.9 gigatons annually. This amount accounts for roughly half of the global yearly carbon emissions from fossil fuel combustion, which is approximately 7.0 gigatons (Elbert et al., 2012). They aggregate the soil, preventing dust storms that might otherwise result in the dissemination of pathogenic bacteria and fungi (Roriguez-Caballero et al. 2022). Biocrusts are known to influence local and regional water cycles as well as plant germination and growth (Weber et al., 2016). Biocrusts are highly susceptible to climate change and are anticipated to experience a 39% total area loss by 2070 (Rodriguez-Caballero et al., 2018). Moss biocrusts are particularly susceptible to climate change; warming has been shown to strikingly decrease moss biocrust cover, without any sign of recovery (Phillips et al., 2022).

It has been suggested that biocrusts are capable of nutrient transfer with vascular desert plants via fungal networks (Rudgers et al., 2018). It is thought that nutrients may be retained in a fungal “loop” by partitioning of resources across space and time within drylands (Fenchel, 2008). The fungal loop hypothesis states that fungi link nitrogen limited plants with nitrogen fixing biocrusts, which differ in their response to rain events as biocrusts contain poikilohydric organisms which require little water to become metabolically active (Ladrón de Guevara and Maestre, 2022). This loop would reduce the loss of nutrients due to leaching, erosion or gaseous loss. The low vegetation density of drylands, especially in hyper arid regions, further highlights the importance of dryland fungi in ecosystem functioning (Weber et al., 2016). Fungi likely dominate biogeochemical cycles in drylands due to their ability to remain metabolically active at much lower soil water potentials than bacteria (Weber et al., 2016).

The multispecies communities that comprise biocrusts are typically extremotolerant and resistant to desiccation (Weber et al., 2022). Fungi are no exception. Previously described fungal constituents of biological soil crusts include Ascomycetes, Dothideomycetes, Agaricomycetes, Chytridiomycetes, Tremellomycetes, Glomeromycetes, Mortierellomycetes, and others (Bates et al., 2012, Zhang et al., 2016, Abed et al., 2019, Pombubpa et al., 2019, Coleine et al., 2024). Black yeasts, which are among the most extreme tolerant Eukaryotes known, are also biocrust constituents (Carr et al., 2023). Black yeasts, or meristematic colonial fungi, are known for their ability to produce melanin, which facilitates their survival from the Atacama Desert, to Antarctica, to the Chernobyl nuclear reactor with high radioactivity (Cantrell et al., 2011, Zakharova et al. 2013, Coleine et al., 2022). Black yeasts are hypothesized to serve as “sunscreen” for biocrusts, blocking harsh UV radiation and shielding other biocrust organisms (Sterflinger et al., 2012, Kurbessoian et al., 2020, Carr et al., 2023). However, black yeasts’ taxonomic composition within biocrusts has not been extensively characterized.

The effects of aridity on the biocrust mycobiome have, similarly, yet to be determined. Warming has been shown to alter fungal community composition in dryland plants, resulting in poor plant performance (León-Sánchez, 2018). In *Agave tequilana,* drought was found to be a key regulator of arbuscular mycorrhizal symbiosis formation. Severe drought was found to inhibit the formation of mycorrhizal symbioses almost entirely (Chávez-González et al., 2024). Similar effects have been documented in grasses (Remke et al., 2021). As fungi have been shown to promote the growth of mosses, a community shift within the moss biocrust mycobiome may partially account for the dramatic losses of moss biocrust cover (Mathieu et al., 2024). Documenting the effects of increased aridity on the moss biocrust mycobiome is crucial to our understanding of climate’s impact on drylands.

To capture the full diversity of fungi associated with moss biocrusts and characterize the impact of aridity on their community composition, a culture-dependent and independent survey was conducted across an aridity gradient in Southern California. This survey encompassed sites across the California Coast as well as the Mojave and Colorado Deserts. We hypothesized that (i) fungal taxa known to be extreme adapted would dominate hyper arid sites; (ii) at sites with lower aridity, there would be higher fungal diversity; (iii) surface sterilization would reveal putatively endophytic taxa. This study provides novel insights into the roles that fungi play in biological soil crusts, as well as how this critical kingdom of life may respond to climate change.

## Methods

### Collection of moss biocrusts

Moss biocrusts were collected across an aridity gradient from the Mojave Desert, Colorado Desert, and California Coast along a Northeastern transect. Each collection site possessed unique features. Cima Volcanic Field (CIMA, 35.1997917 N, 115.8699722 W) in the central Mojave Desert contains volcanic rock which, due to its black color, absorbs solar radiation and makes this site extremely hot. The Granite Mountains (GMT, 34.7804861 N, 115.6297361 W) in the Southern Mojave Desert is characterized by granitic soil and is dominated by both woody shrubs and Joshua Trees (Table S1). Mojave Desert sites in this study were classified as hyper-arid (Table S1).

Oasis De Los Osos Reserve (ODLO, 33.89182 N, 116.68908W) lies in the Northern Colorado Desert. It possesses a mixture of continental desert and Mediterranean climate due to the presence of a perennial water source. Anza Borrego Research Center (AB, 33.2407889 N, 116.3895333 W) is situated in the eastern Colorado Desert on the Northern edge of Anza-Borrego Desert State Park. The Colorado Desert experiences milder temperature swings and greater water availability than the Mojave. These collection sites were classified as arid (Weiss and Overpeck, 2005). Torrey Pines State Preserve, a semi-arid coastal site, is classified as a Mediterranean climate and hosts many rare and endangered species. We collected from two locations in Torrey Pines: one on a vegetation-free sandstone bluff (TP, 32.9456861 N, 117.2494806 W), and another with sandy soil and woody shrubs nearby (TP2, 32.9421506 N, 117.2482757 W). Across all collection sites, moss biocrusts displaying a dark, short morphotype were collected in triplicate from each site (Table S1). To compare the moss mycobiome to the adjacent bare soil, the nearest patch of bare soil was opportunistically sampled in triplicate. A metal spatula sterilized with 70% ethanol between samples was used to collect biological soil crusts or scoop dirt into 50 mL falcon tubes or sterile bags. Samples were stored in a cooler containing dry ice in the field and transported to the University of California, Riverside and stored at −20C.

### Metabarcode sequencing

All metabarcode data from this study are associated with BioProject accession number PRJNA1148194 (Kelly et al. 2024). Moss biocrust samples were divided into three fractions: dirt adhering to the crust, untreated biocrust, and surface-sterilized biocrust. Surface sterilization was carried out via the methodology outlined in U’Ren et al. (2010). The biocrust samples were rinsed using high pressure water to remove excess dirt. They were then subjected to sequential treatments of 95% ethanol for 30 seconds, 10% bleach (0.5% NaOCl) for 2 minutes, and 70% ethanol for 2 minutes. After air-drying in a sterile hood, samples were stored at −20°C.

DNA was extracted from .25 g of bare soil or biocrust samples using the QIAGEN DNeasy PowerSoil kit (Qiagen, Germantown, MD, USA). The manufacturer’s protocol used with only a single modification: bead beating was extended by ten-minutes to enhance DNA extraction from plant tissues. PCR amplification targeted the ITS1 region of fungal ribosomal DNA using ITS1F and ITS2 primers following the Earth Microbiome Project (EMP) guidelines (Thompson et al., 2017). PCR reactions were prepared in a 25 µL reaction volume with 10 µL 2X Platinum Hot Start PCR Master Mix (Thermo Fisher Scientific Inc., Waltham, MA, USA), 1 µL template DNA, 0.5 µL of 10 µM forward and reverse primers, and 13 µL PCR-grade water (Sigma-Aldrich, St Louis, MO, USA). The thermal cycling protocol used a C1000 thermal cycler (BioRad, Hercules, CA, USA) as follows: initial denaturation at 94°C for 1 minute; 35 cycles of 94°C for 30 seconds, 52°C for 30 seconds, and 68°C for 30 seconds; final extension at 68°C for 10 minutes.

PCR products were normalized and purified using the Invitrogen SequalPrep kit (Invitrogen, Waltham, MA, USA). NEBNext Ultra II library prep kit was used to generate libraries that were sequenced on Illumina platforms (MiSeq or NextSeq2000). This generated paired-end reads of 2×300 bp at the University of California, Riverside Institute for Integrative Genome Biology Core (http://iigb.ucr.edu). Although a small subset of samples underwent 2×250 sequencing, all read lengths were trimmed to 250 bases during bioinformatic processing to ensure uniformity.

### Culturing methods

Biocrust associated yeasts were cultured using a soil dilution protocol and multiple media types. 0.5 grams of each biocrust sample was added to a 15 mL falcon tube along with 10 mL of sterile water. The mixture was then incubated in a shaking incubator at 180 rpm and 26.5°C for 24 hours. After incubation, 200 µL of the solution was plated onto Cornmeal agar (CMA) with chloramphenicol (Cm), Rhodotorula medium (RM, 1g MSG/L) with Cm, and low MSG RM (0.1g MSG/L) + Cm. The plates are then left at 18°C. Subculturing of yeast colonies began between two and seven days of growth, focusing on colonies that appeared morphologically distinct. This process was carried out using biocrust collected from AB, TP, GMT, CIMA, and ODLO.

### Genotyping cultures

A molecular barcode useful for fungal species identification was generated by amplifying the ITS1-5.8S-ITS2 region via PCR and Sanger sequencing of the amplicon product (White et al., 1990). DNA was first extracted using either the QIAGEN DNA Blood and Tissue Kit (Qiagen, Germantown, MD, USA) or prepman ultra kit (Applied Biosystems, Waltham, MA, USA). The PCR protocol for ITS1 (5’-TCCGTAGGTGAACCTGCGG-3’) and ITS4 (5’-TCCTCCGCTTATTGATATGC-3’) followed standard procedures: 94°C for 30 seconds, then 30 cycles of 94°C for 30 seconds, 50°C for 60 seconds, 68°C for 60 seconds. A final extension step of 68°C for 5 minutes concluded the run. A biological safety cabinet was sterilized under UV light with 70% ethanol. Reaction mixtures were prepared as follows: OneTaq® Quick-Load® 2X.

Master Mix with Standard Buffer (New England Biolabs, Ipswich, MA, USA), .5 µL of 10 µM forward and reverse primer, 3 µL template DNA, and 8.5 µL nuclease-free water (Thermofisher Scientific, Kalamazoo, MI, USA). PCR products were stored at −20°C and analyzed by gel electrophoresis. The Macherey-Nagel NucleoSpin Gel and PCR Clean-up kit (Düren, Nordrhein-Westfalen, Germany) was used to purify PCR products before sanger sequencing at the Institute for Integrative Genome Biology, Core Facilities, University of California Riverside (http://iigb.ucr.edu). Cultures were identified by sequence search against the UNITE v9.3 database (Abarenkov et al., 2024) using the AMPtk default hybrid approach.

### Bioinformatics and Data Analysis

Demultiplexed metabarcode samples underwent processing and analysis using AMPtk version 1.60 (Palmer et al., 2018) (https://github.com/nextgenusfs/amptk). VSEARCH was used to merge paired-end reads (version 2.28.1) (Rognes et al., 2016). A mismatch of up to two base pairs was allowed for primer removal, and sequences were trimmed to a length of 250 base pairs. Reads shorter than 100 bp were discarded. Following preprocessing, 6,972,409 valid output reads remained. Quality filtering was applied with an expected error parameter of 1.0, resulting in 6,056,727 reads retained for further analysis. The DADA2 pipeline was employed to cluster these reads into Amplicon Sequence Variants (ASVs) (Callahan et al., 2016). After removing a total of 1051 de novo and reference chimeras using VSEARCH (version 2.28.1) and validating orientations, 10,413 ASVs were identified (Rognes et al., 2016).

Taxonomic assignments were performed by searching against the UNITE v9.3 database (Abarenkov et al., 2024) using the AMPtk default hybrid approach, which employs a Global Alignment hybrid taxonomy assignment to determine consensus last common ancestors. This method prioritizes the most specific taxonomic classification in cases of conflicting results (Palmer et al., 2018). Subsequently, multiple sequence alignment across all ASVs was conducted using MAFFT (v7), and a phylogenetic tree was constructed from this alignment using FastTree (v2.1) (Katoh and Standley, 2013; Price et al., 2010). For the comparison of ASVs, culture ITS sequence, and NCBI type species, an alignment was constructed using MAFFT (v7), and the alignment was trimmed using ClipKIT (2.4.1) (Katoh and Standley, 2013, Steenwyk et al., 2020) A phylogenetic tree was constructed from this alignment using IQtree (v2.4) (Nguyen et al. 2015).

### Statistical Data Analysis and Visualization

Data analyses were conducted using R version 4.4 (R Core Team, 2024), RStudio version 1.4.1103-4 (RStudio Team, 2023), and Phyloseq (Xu et al., 2022; McMurdie and Holmes, 2013). All samples were rarefied to a depth of 7,800 reads, which resulted in the exclusion of one sample. Differences in alpha diversity were assessed using ANOVA and Tukey’s HSD test implemented in base R. Differences in beta diversity calculated using Bray-Curtis dissimilarity were evaluated using a PERMANOVA test with the ’adonis’ function from the vegan package version 2.6-7 in R (Oksanen et al., 2024). Venn diagrams were constructed using the ’ps_venn’ function from MicoEco v0.9.15 (Russel, 2021). Phylogenies were visualized using the ggtree package (Xu et al., 2022). Differences in relative abundance were determined using DEseq using methodology described by McMurdie and Holmes, 2014 (Love et al., 2014).

### Microscopy

Fungal colonization of moss tissue was visualized via a modified version of the Hanke and Rensing (2010) protocol. Moss tissue from each site, stored at −20°C, was carefully washed with water. Samples were subjected to the following treatment: 10% KOH at 95°C for ten minutes, 3 minutes at room temperature in 5% HCl. Samples were stained using 0.1% lactophenol blue for twenty minutes at room temperature, and de-stained using 50% glycerol for 24 hours. They were then placed on a slide with glycerol and observed using an OMAX M83EZ Series Trinocular Compound Microscope (AMscope, Irvine, CA, USA).

## Results

### Local climate influences fungal diversity

Beta diversity was significantly different between local climates (*p* < .01) (Fig. 2A). A trend of increased inter-sample beta diversity homogeneity with increased site aridity was observed (Table S1). There was also a signature of community similarity by site (Fig. S1).

**Figure 1.**
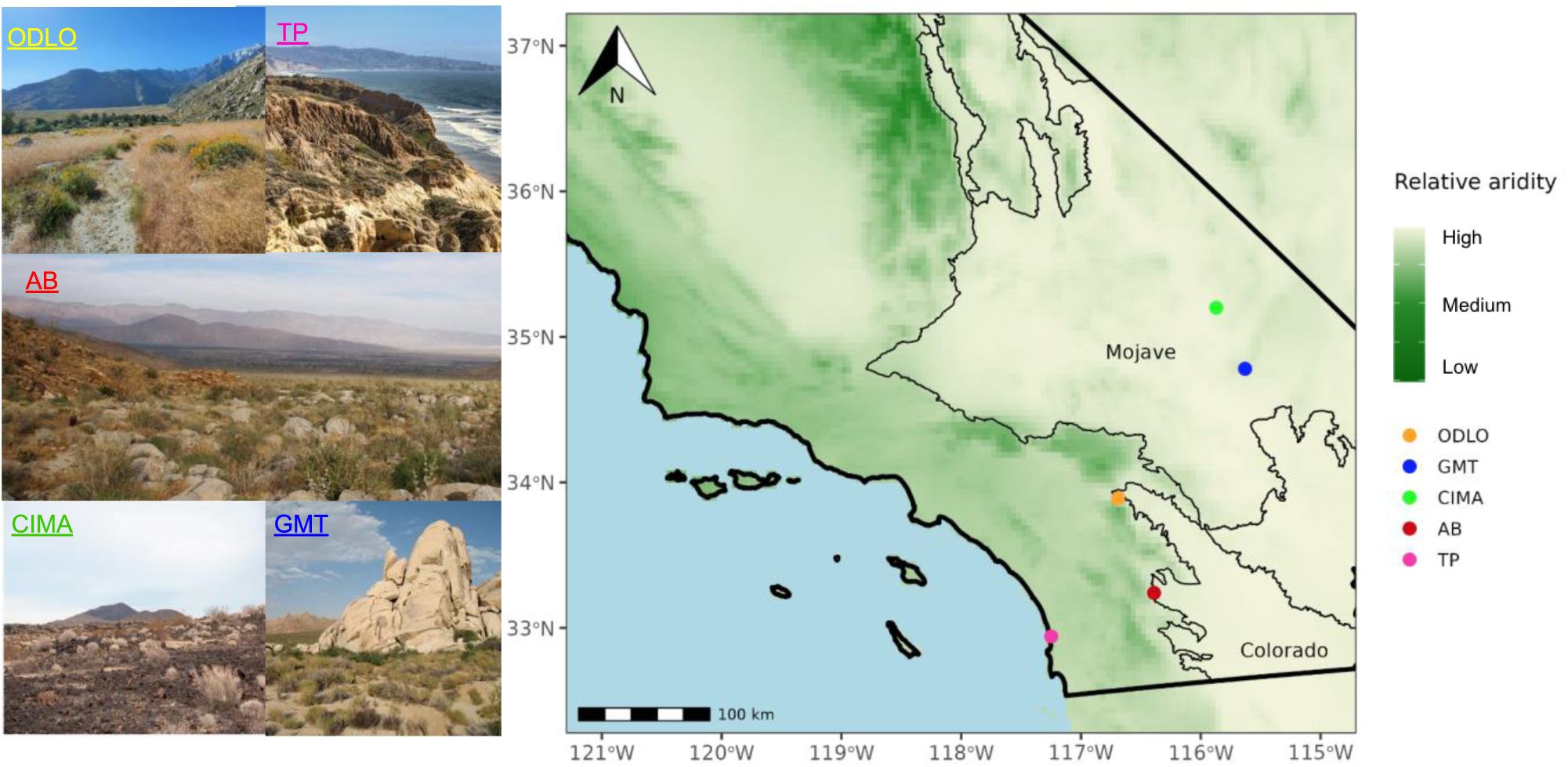
Field sampling sites. Moss biocrusts were collected along an aridity gradient in a Northeast transect. Aridity index was obtained from the Terraclimate dataset. ODLO = Oasis De Los Osos Reserve, TP = Torrey Pines, AB = Anza Borrego Research Station, CIMA = CIMA Volcanic Field, GMT = Granite Mountains Research Center. This figure was adapted from Kelly et al. 2024.

**Figure 2.**
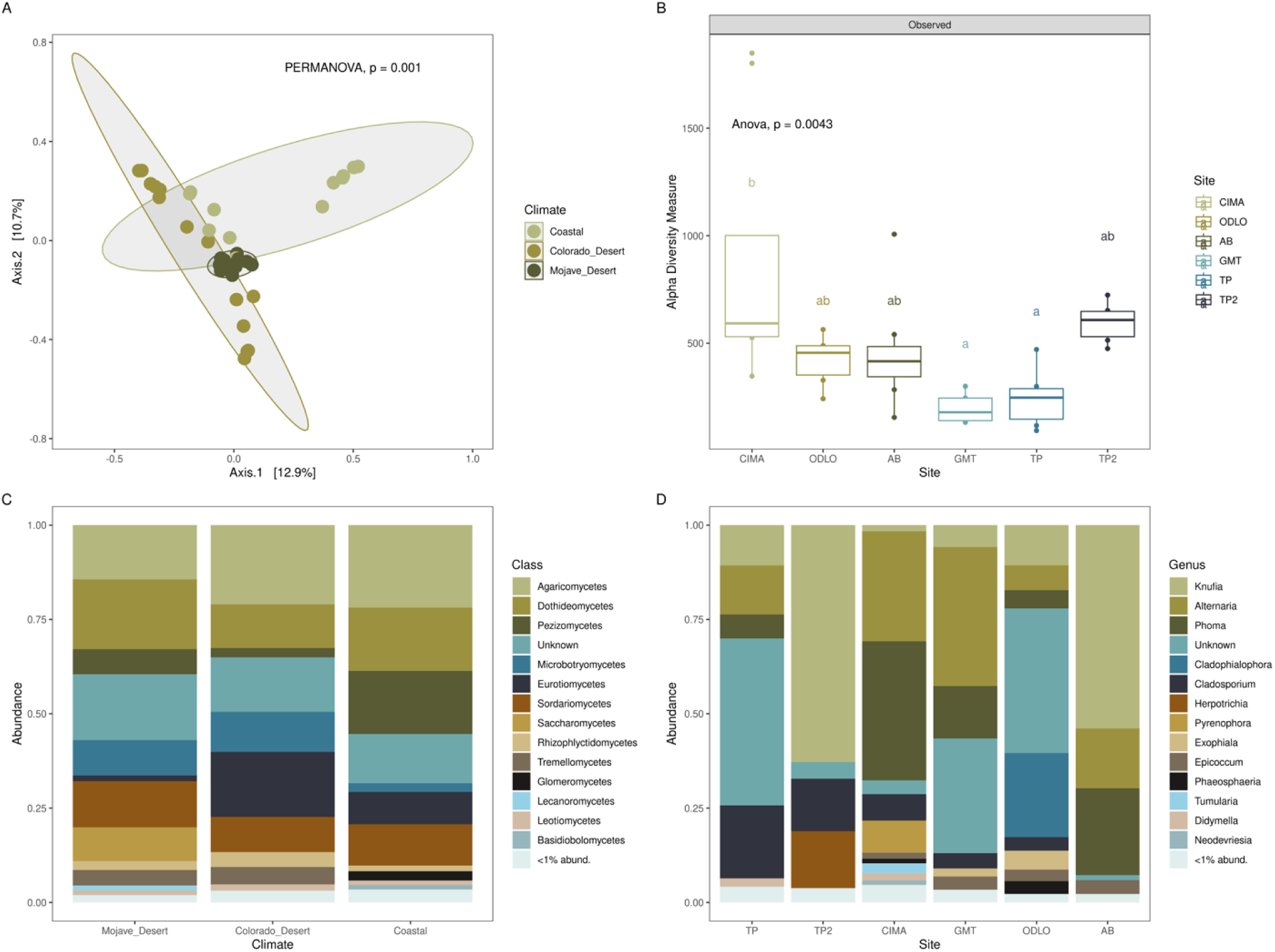
(A) Beta diversity of the moss biocrust mycobiome by climate. (B) Alpha diversity of the moss biocrust mycobiome by site. (C) Taxonomic composition of moss biocrust mycobiome by climate. (D) Black yeast taxonomic composition by site.

A nuanced relationship between alpha diversity and aridity was apparent. Notably, CIMA exhibited a significantly higher alpha diversity of fungi than all other sites except for TP2 (*p* < .01) (Fig. 2B). This is remarkable as CIMA is a hyper-arid site (Table S1). At TP, alpha diversity was approximately as low as GMT despite the fact that TP is semi-arid and GMT is hyper-arid. TP2 possessed the highest alpha diversity overall.

### Taxonomic Composition of Biocrust Fungi Shifts with Local Climate

Fungal community composition varied significantly across local climates with differing aridity indices (Fig. 2C). Taxonomic variation in the biocrust mycobiome was substantial between sites, even at the class level (Fig. S3). Semi-arid coastal sites showed a markedly higher relative abundance of Glomeromycetes (Mucoromycota), Pezizomycetes (Ascomycota), and Basidiobolomycetes (Basidiomycota) compared to the arid and hyper-arid regions of the

Mojave and Colorado Deserts. In contrast, Microbotryomycetes (Basidiomycota) were more abundant in the Mojave and Colorado Deserts than in coastal sites. Notably, the Colorado Desert sites were dominated by Agaricomycetes (Basidiomycota), with Agaricomycetes comprising approximately 75% of fungal reads at ODLO (Fig. S2). Rhizophylictidomycetes (Chytridiomycota) were present across all climate types, while Saccharomycetes were enriched specifically in the Mojave Desert (Fig. 2C).

Unclassified taxa represented a significant proportion of fungal reads at some sites, comprising over 25% at AB (Fig. S2). Several fungal taxa were more abundant in semi-arid climates, including *Glomus macrocarpum*, which showed higher relative abundance in these regions (Fig. 3, Table S2). Black yeast composition also varied significantly at the genus level between sites (Fig. 2D). For instance, *Knufia sp.* (Ascomycota; Chaetothyriales) dominated at TP2 and AB, whereas *Cladophialophora sp.* (Ascomycota; Chaetothyriales) was prevalent at TP and ODLO. In contrast, *Phoma sp*. (Ascomycota; Dothideomycetes) was the dominant black yeast at CIMA and GMT. A large proportion of black yeast taxa, nearly 50% at TP and ODLO, could not be classified to even the genus level. Black yeast species such as *Exophiala heteromorpha*, *Knufia sp.*, *Cryptococcus neoformans*, and *Cladophialophora sp.* were enriched in arid and hyper-arid climates (Fig. 3, Table S2). Other non-melanized fungi including *Naganishia vishniacii* were also enriched in arid and hyper-arid environments. Several fungal ASVs showed significant differential abundance between the Mojave and Colorado Desert (Fig. S7)

**Figure 3.**
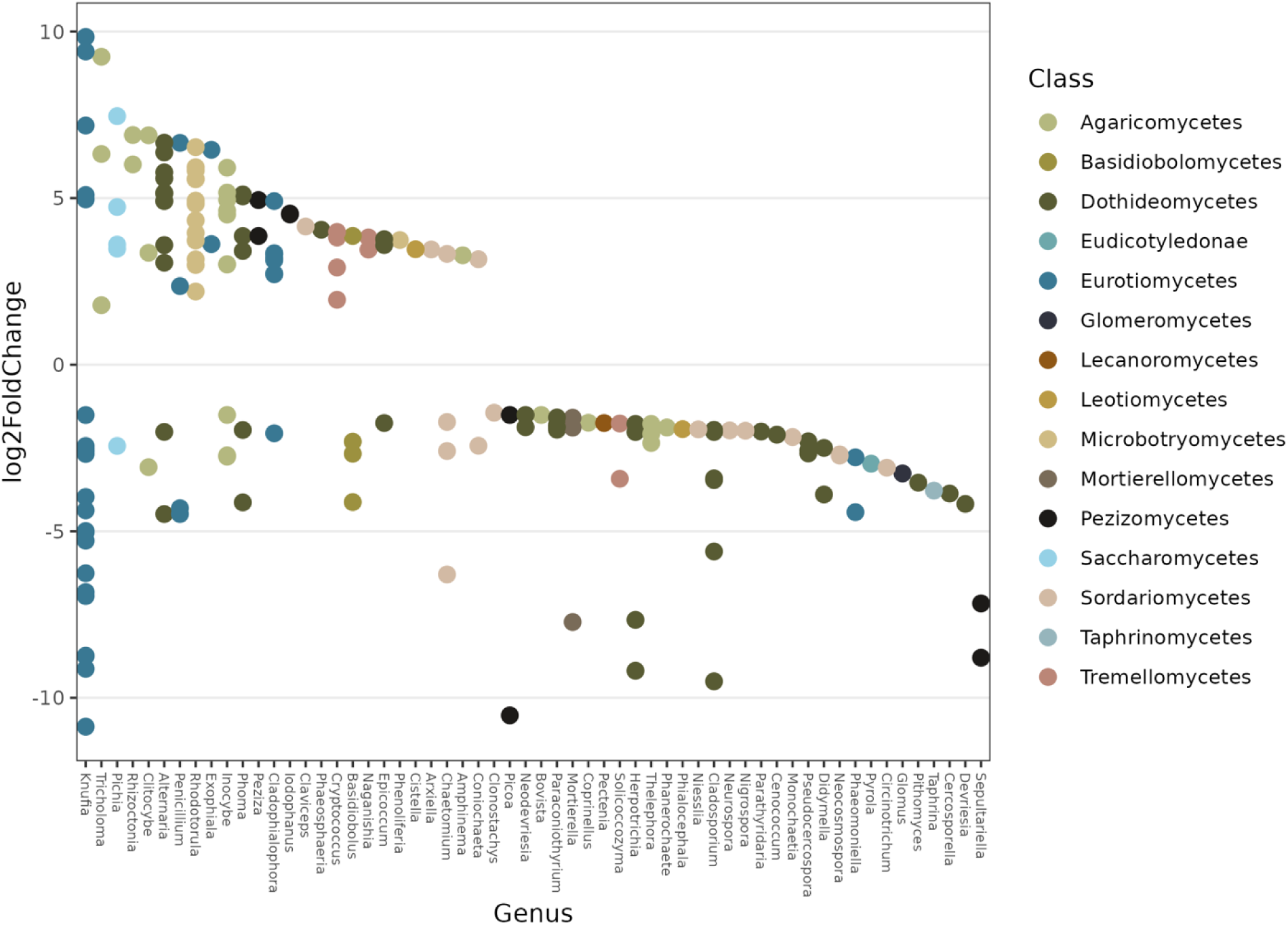
Significant relative abundance changes in fungal taxa in arid and hyper arid regions compared to semi-arid regions calculated using DEseq2.

### Culture-dependent survey reveals a dominance of putatively extreme tolerant taxa in dryland biocrusts

A total of 75 isolates were cultured from moss crusts and lichen crusts. A comprehensive list of isolated fungal taxa and first reports is provided in Table S3. Overall, Tremellomycetes including *Naganishia spp.* and *F. magnum* were the dominant fungal taxa isolated from moss biocrusts. Dothideomycetes (e.g *Myriangium duriaei*) followed by Tremellomycetes (e.g *Papillotrema terrestris*) were the most dominant fungal classes within lichen crusts (Fig 4A).

**Figure 4.**
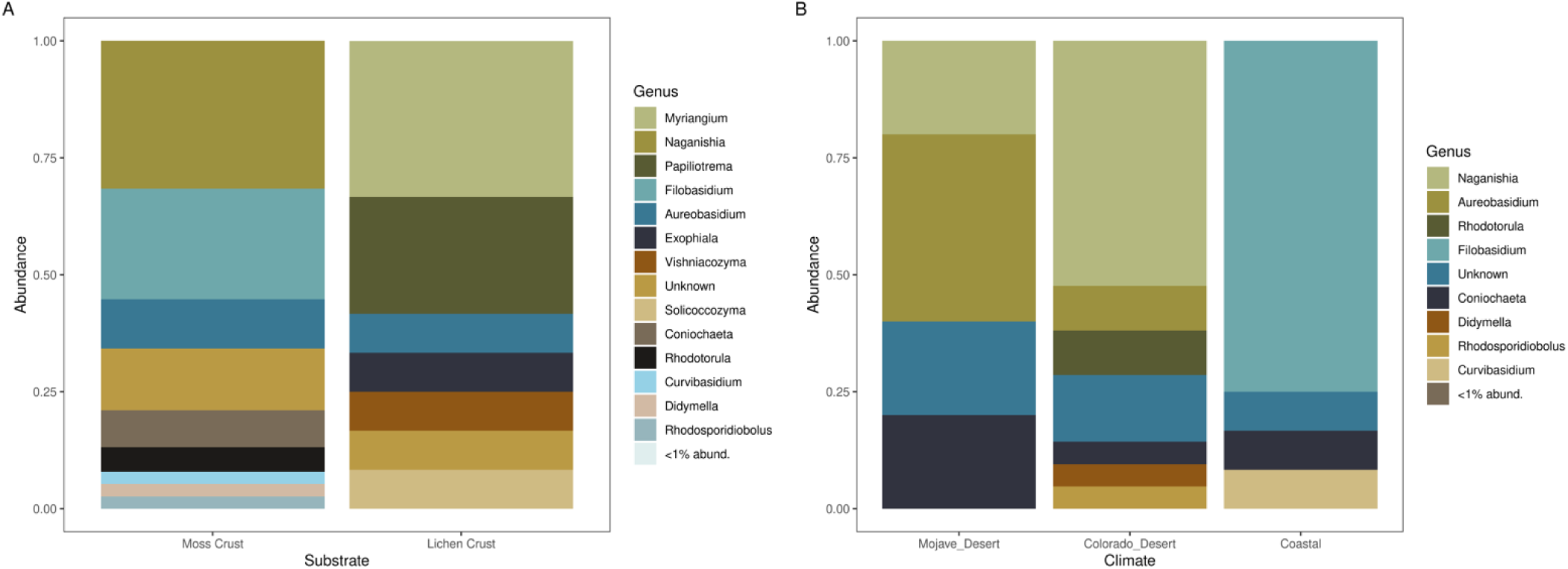
Results of culture-dependent survey. (A) Taxonomic identities of fungal isolates from moss biocrusts (58 isolates) compared to lichen dominated biocrusts (15 isolates). (B) Taxonomic identities of fungal cultures associated with Mojave Desert (24 isolates), Colorado Desert (22 isolates), and Coastal moss crusts (12 isolates).

Isolates of *Rhodotorula kratochvilovae* and *Coniochaeta boothii* were found only from moss biocrusts (Fig 4A, Table S4). In addition, several cultured taxa from both moss and lichen biocrusts could not be classified to the genus level with ITS sequence. Moss crust from the coast was dominated by *Filobasidium magnum* (Tremellomycetes) (Fig 4B, Table S4). Yeasts in the genus *Naganishia* (Tremellomycetes) were the most abundant genera within the Colorado Desert, including species *N. globosa*, *N. onofrii* and *N. diffluens* (Fig 4B, Table S4).

*Aureobasidium* (Dothideomycetes) was the dominant genus in the Mojave Desert, including species *A. pullans and A. melanogenum*. The genus *Coniochaeta* (Sordariomycetes) including *C. boothii*, *C. luteorubra*, and *C. polymorpha* (Sordariomycetes) was recovered from the Colorado Desert, Mojave Desert, and the coast respectively. In the culture-dependent survey, only two *Exophiala* isolates were recovered, whereas the culture-independent survey identified a total of 39 *Exophiala* ASVs, most of which clustered distinctly from NCBI type species (Fig 5).

**Figure 5.**
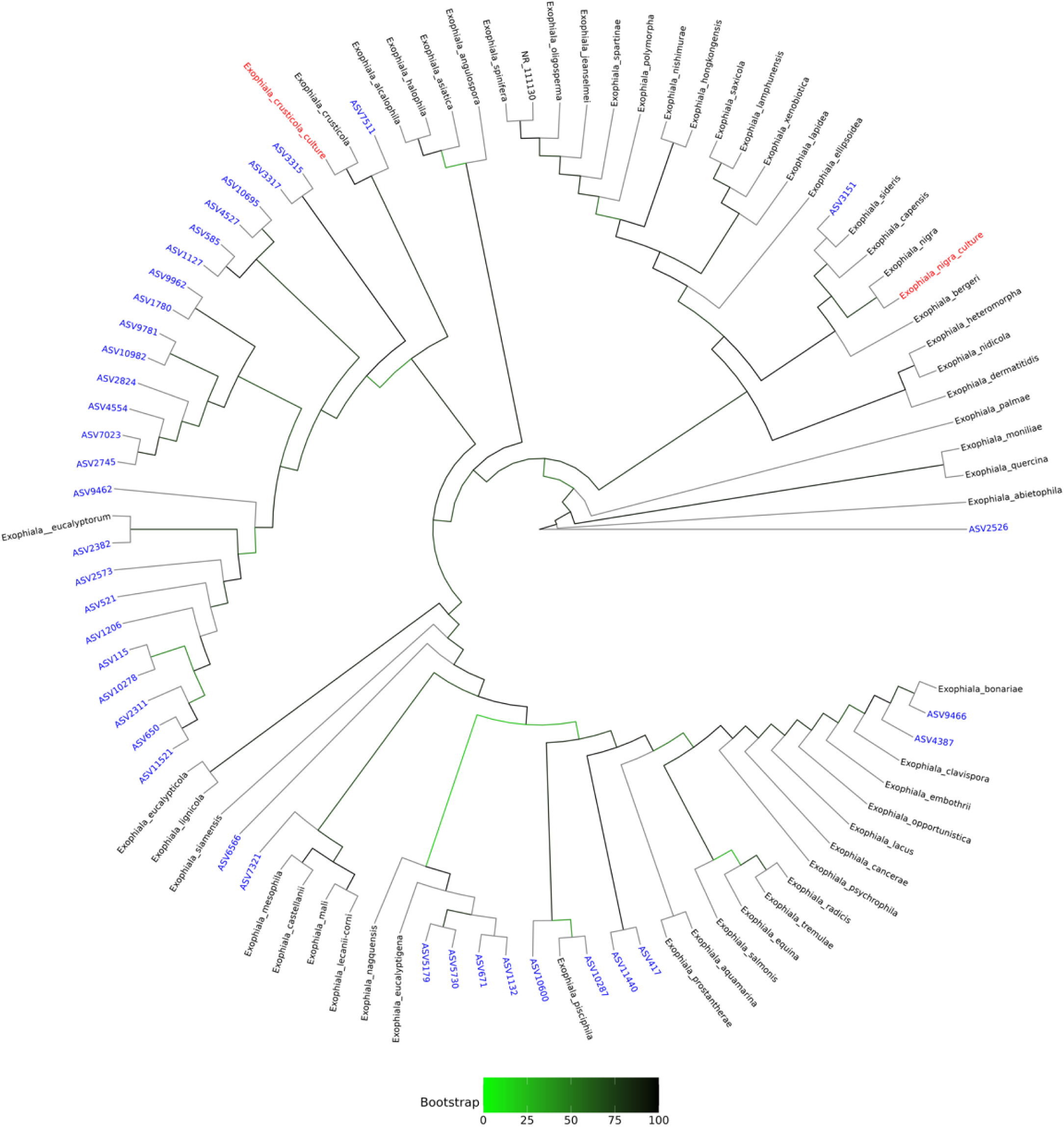
Cladogram of *Exophiala* ASVs from the culture-independent survey (blue), isolates obtained from the culture-dependent survey (red), and *Exophiala* NCBI type species (black). Low bootstrap values are green while high values are black. Many ASVs cluster distinctly from NCBI type species while isolates more closely resemble known species.

### Endophytic and epiphytic taxa associated with moss biocrusts

At the class level, the taxonomic composition across substrates (i.e., moss crust, sterilized moss, bare soil, and soil attached to the crust) was largely consistent (Fig. S3A). However, surface sterilization markedly increased the relative abundance of Microbotryomycetes. Specifically, *Rhodotorula mucilaginosa* and *R. paludigena* were enriched in sterilized moss crust samples (Table S4, Fig. S3C).

Surface sterilization uncovered approximately 850 unique ASVs that would have otherwise not been detected (Fig. S3B). A complete list of ASVs with significantly higher relative abundance following surface sterilization is provided in Table S3. Black yeast genera such as *Knufia* and *Exophiala* were notably reduced in abundance after sterilization, suggesting these fungi primarily reside on the moss surface (Fig. S3C). *Neodiversia* was present both before and after sterilization. *Knufia* was dominant in biocrust-associated samples but maintained relative abundance below 1% in bare soil (Fig. S3D).

Moss biocrust fungal beta diversity was influenced by surface sterilization (Fig. S4). Sterilized moss crusts exhibited greater inter-sample community similarity compared to unsterilized moss in terms of beta diversity. In terms of alpha diversity, there was no difference before and after surface sterilization (Fig. S5). Comparing bare soil to moss crust, *Cr. neoformans*, *R. mucilaginosa*, and *G. macrocarpum* were significantly enriched in moss crust samples (Table S5). Microscopy further revealed colonization of healthy moss tissue by septate fungi in *Crossidium squamiferum* at site AB (Fig. S6), supporting the role of biocrust fungi as endophytes of moss. Several taxa including *Rhodotorula* spp. were enriched in moss biocrusts compared to adjacent bare soil (Fig. S8)

## Discussion

Our study provides insights into: (i) the impact of local aridity on fungal community diversity and composition; (ii) endophytic composition of biocrust mosses; (iii) the diversity of black yeast and unclassified fungal taxa associated with moss biocrusts

### Fungal community similarity and composition are putatively influenced by the local aridity index

Previous studies have shown that fungal diversity and abundance are reduced at higher aridity in soils (Maestre et al., 2015). However, the effects of aridity on fungal diversity and composition in biocrusts remain understudied. Fungi play critical roles in biocrusts, including nutrient cycling and soil aggregation (Bates et al., 2006). Shifts in fungal diversity driven by climate change are likely to significantly impact biocrust and dryland ecology (Zhang et al., 2024). This study found increased fungal beta diversity homogeneity at higher aridity, which may help explain observed losses of biocrust cover under warming scenarios (Phillips et al., 2022). Taken together, these results suggest that fungal communities are strongly influenced by aridity, with implications for the ecosystem services provided by biocrusts. These findings are important considerations for land managers, who may need to prioritize conservation efforts for biocrusts in drylands under climate change.

In arid to hyper-arid climates, biocrust fungal communities shifted toward extreme-tolerant taxa, including *Rhodotorula* spp. (Microbotryomycetes). Members of the *Rhodotorula* genus are polyextremotolerant and widely distributed in drylands globally (Duarte et al., 2013, Selbman et al., 2014, Wei et al., 2022). For example, *R. mucilaginosa*, often isolated in Antarctic environments in addition to being a cosmopolitan and widely abundant species, was enriched in arid to hyper-arid climates in this study (Butinar et al., 2007, Wei et al., 2022). Many of the cultured fungi were pigmented, producing melanin or carotenoids, which are known to enhance resistance to UV radiation, desiccation, and oxidative stress; key adaptations for survival in drylands (Coleine et al., 2022).

Basidiomycetous yeasts were also enriched in hyper-arid sites. *Rhodotorula kratochvilovae* cultures were obtained from multiple arid sites, indicating its putative viability in these environments. *Cryptococcus neoformans* and *Naganishia vishniacii* (Tremellomycetes) were also more abundant in arid sites compared to semi-arid sites. These yeasts are frequently isolated from soils in extreme environments (Butinar et al., 2007, Buzzini et al., 2018) and have been shown to play critical roles in soil carbon stabilization, as seen in Arctic ecosystems (Trejos-Espeleta et al., 2019). The ecological roles of basidiomycetous yeasts in biocrusts remain poorly understood, but their abundance and diversity in arid climates highlight their potential importance in dryland soil processes.

Severe drought has been shown to inhibit mycorrhizal symbiosis formation in plants like *Agave tequiliana* (Chávez-González et al., 2024). Similar effects have been documented in grasses, with the loss of symbiosis associated with reduced plant growth (Remke et al., 2021). As fungi are known to promote moss growth, reduced fungal diversity may partly explain the dramatic losses in moss biocrust cover under warming scenarios (Mathieu et al., 2024). Interestingly, species richness was not consistently lower in hyper-arid sites; in some cases, alpha diversity was higher, reflecting an abundance of putatively niche-specialized taxa. These findings align with previous work highlighting fungal specialization in drylands (Selbman et al., 2005, Sterflinger et al., 2012, Talbot et al., 2014).

Site strongly influenced biocrust mycobiome beta diversity, corroborating previous findings of biogeographical patterns in fungal community composition (Pombubpa et al., 2020). Future studies should employ extensive sampling across drylands at continental or global scales to determine whether these patterns are consistent beyond California drylands. This could include combining metabarcoding with culturing to better characterize fungal diversity and functional roles in biocrust ecosystems.

### Fungi act as endophytes and epiphytes of biocrust mosses

Surface sterilization of healthy moss biocrusts revealed black yeast including *Cladophialophora* sp., suggesting that these fungi may act as endophytes of desert moss. *Rhodotorula mucilaginosa* and *R. paludigena* were also significantly enriched after surface sterilization of biocrust mosses. This aligns with their known roles as plant endophytes (Khunnamwong et al., 2018, Peng et al., 2018, Sen et al., 2019).

*Rhodotorula paludigena* has been shown to produce Indole Acetic Acid (IAA), a key plant growth hormone (Peng et al., 2018). Surface sterilization also reduced the abundance of possible epiphyte ASVs, including *Exophiala heteromorpha*, which are melanized black yeast. These yeasts may contribute to protecting moss from harsh UV radiation in drylands.

Many taxa were enriched in moss biocrust associated samples compared to adjacent bare soil and soil clinging to the moss biocrust. These included *Sporobolomyces roseus*, *Rhodotorula mucilaginosa*, *Glomus macrocarpum*, *Cr. neoformans*, and other species that are potentially plant endophytic (Kapoor et al., 2002, Sen et al., 2019, Sun et al., 2023). *Coniochaeta* spp. were also frequently isolated from moss biocrusts, though they were often not identified to the species level. Some *Coniochaeta* species have been documented as moss endophytes and shown to promote moss growth, but the genus has not been previously reported in the Mojave or Colorado Deserts (Arnold et al., 2021, Mathieu et al., 2024). *Aureobasidium pullulans*, another common endophyte and black yeast, was also recovered in the culture-dependent survey from moss biocrust (Ali et al., 2019).

Intracellular colonization within healthy plant cells is a diagnostic feature of symbiosis, but most mosses are thought to be asymbiotic (Field et al., 2015). Recent research has demonstrated that fungi can promote moss growth (Mathieu et al., 2024). In this study, the visualization of septate hyphal colonization within healthy moss tissue suggests that fungi in biocrusts may act as moss endophytes. This highlights the potential for biocrust fungi to support the health of dominant photoautotrophs. Further research is needed to better understand fungal-bryophyte interactions in biocrusts and whether fungi can enhance biocrust moss growth.

As we face a warmer, drier climate, gaining a deeper understanding of desert symbioses will facilitate the use of plant symbiotic fungi as inoculants to improve stress resistance in crops. Our culture-dependent and independent surveys revealed that *Rhodotorula* species are enriched in healthy moss tissue after surface sterilization and in mosses from arid to hyper-arid climates compared to coastal mosses. *Rhodotorula* species have also been shown to promote stress tolerance in plants (Silambarasan et al., 2019). Based on these findings and their known roles as plant endophytes mediating stress resistance, *Rhodotorula* species could potentially be used as bioinoculants to enhance moss biocrust resistance to climate change. Since moss biocrusts are among the most vulnerable biocrust types to climate change, such inoculants could provide critical protection against its effects. Further work is needed to determine if *Rhodotorula* spp. can promote moss biocrust growth and stress resistance.

### Diverse and undescribed fungi are rich within biocrusts

Many fungi identified in this survey could not be assigned taxonomy even at the class level. For black yeasts, nearly 50% of total reads at some sites could not be assigned taxonomy to the genus level. Similarly, a large proportion of cultured taxa from both moss and lichen biocrusts could not be classified to the genus or species level. These findings indicate that biocrusts harbor diverse and undescribed fungi that should be prioritized in taxonomic studies. This aligns with previous research suggesting that extreme environments are ideal for addressing gaps in the Eukaryotic tree of life (Rappaport et al. 2023). Targeting these fungi could lead to the identification of numerous undescribed species and provide insights into their roles in ecosystem functioning and eukaryotic genomic adaptations to hyper-arid climates. There is a lack of studies focusing on the mycobiome of the Mojave Desert, with the Northern, Eastern, and Western Mojave regions remaining poorly sampled (Pombubpa et al., 2020). Future efforts should prioritize comprehensive, region-wide sampling paired with metagenomic and culturomic approaches to systematically document and characterize the understudied fungal communities in the Mojave Desert or other extreme environments. Interestingly, Rhizophylictidomycetes (Chytridiomycota) were present in moss biocrusts across all climates. This is consistent with previous studies identifying members of Chytridiomycota in biocrusts (García-Carmona et al., 2022). Further work is needed to determine their trophic mode within moss biocrusts.

The culture-independent survey proved to be more sensitive than the culture-dependent survey for most fungal taxa. For instance, the culture-independent survey revealed 39 taxa of *Exophiala* that were not detected in the culture-dependent survey, while only one cultured isolate was distinct from any amplicon sequence variant (ASV). This is likely due to the unique growth requirements of black yeasts, which are typically slow-growing and often outcompeted by faster-growing fungi (Sterflinger et al., 2012).

Additionally, many fungi recovered from the culture-dependent survey could not be classified to the genus level, emphasizing that culturing fungi from biocrusts often yields undescribed taxa. Culturing also yielded isolates not previously reported at field sites, including the first record of the *Naganishia* genus in the Mojave Desert, *Rhodotorula kratochvilovae* in the Colorado Desert, and *Coniochaeta spp*. in the Mojave.

Both culture-dependent and culture-independent approaches identified numerous black yeasts, including *Knufia* spp., *Aureobasidium* spp., *Exophiala* spp., *Phoma* spp., and *Cladophialophora* spp., among others. Polyextremotolerant *Exophiala* species including *Exophiala viscosa* have been described in biocrusts and have been shown to excrete copious melanin (Carr et al., 2020). This melanin may provide a critical service to biocrust communities: protection from UV radiation. It has also been hypothesized that *Exophiala viscosa* has lichen-like qualities, interacting with cyanobacteria, algae, and other members of the biocrust community to form symbioses (Carr et al., 2020). One isolate from our study*, Exophiala crusticola*, is notable for producing exopolysaccharides that likely contribute to biocrust aggregation and erosion prevention (Bates et al. 2006). It’s precise ecological role within biological soil crusts remains unknown. This is the first study to comprehensively survey the composition of black yeasts in biological soil crusts using a culture-independent approach. Future studies integrating both culture-dependent and culture-independent surveys focusing on black yeasts should be conducted in under sampled regions of the Mojave and Sonoran Deserts to further characterize their diversity and ecological roles in these environments.

### Conclusions

In summary, fungal beta diversity increased in inter-sample homogeneity with increasing aridity. Fungal taxonomic composition shifted toward a higher abundance of extreme adapted fungi at high aridity. These results highlight key factors for land managers to consider, emphasizing the need to prioritize biocrust conservation in drylands facing climate change. Fungal species alpha diversity was influenced by the site. The high alpha diversity of fungi at hyper-arid sites suggested that there may be a variety of niche adapted fungal species within hyper arid sites. Fungi within moss biocrusts appeared to act as endophytes, based on intracellular colonization of the mosses by fungal hyphae within healthy plants. Surface sterilization revealed known fungal taxa within mosses that are common endophytes of vascular plants. Specific taxa including *Rhodotorula spp*. known to promote plant stress resistance were enriched in mosses after surface sterilization. Hence, the survey indicates fungal taxa that may be subjected to further investigation as bioinoculants to promote biocrust health and promote resilience to climate change.

Unsurprisingly, the culture-independent survey of our study was more sensitive than the culture-dependent survey. Still, the culture-dependent survey recovered a handful of taxa not found in the culture-independent survey. Many of the fungi within biocrusts were not able to be assigned taxonomy to the class level, indicating that biocrusts are an ideal sampling target to deepen our understanding of fungal taxonomy. As much as half the total black yeast diversity at some sites, along with many of the culturable fungi, was unable to be assigned taxonomy to the genus level. This indicates biocrusts are a prime sampling target to fill in the unknown lineages of fungi. Future studies should conduct extensive sampling of under-sampled regions of the Mojave Desert to obtain cultures of biocrust associated black yeast. This would lead to a deeper understanding of the taxonomy of biocrust fungi as well as their genomic adaptations to extreme climates.

## Acknowledgements

We are grateful to the Mojave National Preserve (Permit MOJA-2023-SCI-0043), as well as the University of California Natural Reserves, including the Anza Borrego Desert Research Center (Application #52877) and Oasis De Los Osos (Application #52886), for granting permission to conduct this research. We also appreciate the support from Torrey Pines State Natural Reserve (permit application number 23-630-03). Special thanks go to Cassie Ettinger, Mark Yacoub, and Cheng-Hung Tsai for their valuable feedback and suggestions. We also acknowledge Jessica Wu-Woods, Leila Shadmani, and Sadikshya Sharma for their assistance with laboratory logistics and management. Additionally, we recognize the use of Creative Commons licensed images: “Vista of Anza Borrego” by John Fowler (CC BY-SA 3.0, cropped), “View from Guy Fleming Trail” by Remember to Breathe (CC BY-NC-ND 2.0, cropped), and “Granite Mountains” by an unknown creator (Public Domain Mark 1.0 Universal). Computational analyses were conducted using the UC Riverside High Performance Computing Cluster, supported by grants from the National Science Foundation (DBI-1429826 & DBI-2215705) and the National Institutes of Health (NIH) (S10-OD016290). JES is a CIFAR fellow in the *Fungal Kingdom: Threats and Opportunities* program and received support from the USDA National Institute of Food and Agriculture Hatch project CA-R-PPA-211-5062-H. KHK is an NSF ExFAB BioFoundry fellow (UEI: G9QBQDH39DF4) and was partially supported by NSF EF-2125066 to JES. Jared Croyle was an NSF REU participant and was funded by NSF-2051131.

## Author Contributions

The study was designed and planned by JES and KHK. JES, XZL, and KHK provided mentorship throughout the project. Field sampling was executed by KHK. DNA extraction, PCR, normalization, and library preparation for metabarcoding were conducted by KHK. Cultures were isolated and genotyped by KHK and JC. Bioinformatics analysis was carried out by KHK and JES. Statistical analysis and data visualization were performed by KHK. JES oversaw project administration and secured funding. KHK drafted and revised the manuscript with input from JES and XZL. All authors reviewed and approved the final version of the manuscript.

## References

Abarenkov, K., Nilsson, R.H., Larsson, K.-H., Taylor, A.F.S., May, T.W., Frøslev, T.G., Pawlowska, J., Lindahl, B., Põldmaa, K., Truong, C., Vu, D., Hosoya, T., Niskanen, T., Piirmann, T., Ivanov, F., Zirk, A., Peterson, M., Cheeke, T.E., Ishigami, Y., Jansson, A.T., Jeppesen, T.S., Kristiansson, E., Mikryukov, V., Miller, J.T., Oono, R., Ossandon, F.J., Paupério, J., Saar, I., Schigel, D., Suija, A., Tedersoo, L., Kõljalg, U., 2024. The UNITE database for molecular identification and taxonomic communication of fungi and other eukaryotes: sequences, taxa and classifications reconsidered. Nucleic Acids Research 52, D791–D797. 10.1093/nar/gkad1039

Abatzoglou, J.T., Dobrowski, S.Z., Parks, S.A., Hegewisch, K.C., 2018. TerraClimate, a high-resolution global dataset of monthly climate and climatic water balance from 1958–2015. Sci Data 5, 170191. 10.1038/sdata.2017.191

Abed, R.M.M., Tamm, A., Hassenrück, C., Al-Rawahi, A.N., Rodríguez-Caballero, E., Fiedler, S., Maier, S., Weber, B., 2019. Habitat-dependent composition of bacterial and fungal communities in biological soil crusts from Oman. Sci Rep 9, 6468. 10.1038/s41598-019-42911-6

Ali, A., Bilal, S., Khan, A.L., Mabood, F., Al-Harrasi, A., Lee, I.-J., 2019. Endophytic Aureobasidium pullulans BSS6 assisted developments in phytoremediation potentials of Cucumis sativus under Cd and Pb stress. Journal of Plant Interactions 14, 303–313. 10.1080/17429145.2019.1633428

Arnold, A.E., Harrington, A.H., Huang, Y.-L., U’Ren, J.M., Massimo, N.C., Knight-Connoni, V., Inderbitzin, P., 2021. Coniochaeta elegans sp. nov., Coniochaeta montana sp. nov. and Coniochaeta nivea sp. nov., three new species of endophytes with distinctive morphology and functional traits. International Journal of Systematic and Evolutionary Microbiology. 10.1099/ijsem.0.005003

Bai, Y., Scott, T.A., Min, Q., 2014. Climate change implications of soil temperature in the Mojave Desert, USA. Front. Earth Sci. 8, 302–308. 10.1007/s11707-013-0398-3

Bates, S.T., Nash, T.H., Garcia-Pichel, F., 2012. Patterns of diversity for fungal assemblages of biological soil crusts from the southwestern United States. Mycologia 104, 353–361. 10.3852/11-232

Bates, S.T., Reddy, G.S.N., Garcia-Pichel, F., 2006. Exophiala crusticola anam. nov. (affinity Herpotrichiellaceae), a novel black yeast from biological soil crusts in the Western United States. International Journal of Systematic and Evolutionary Microbiology 56, 2697–2702. 10.1099/ijs.0.64332-0

Berdugo, M., Delgado-Baquerizo, M., Soliveres, S., Hernández-Clemente, R., Zhao, Y., Gaitán, J.J., Gross, N., Saiz, H., Maire, V., Lehmann, A., Rillig, M.C., Solé, R.V., Maestre, F.T., 2020. Global ecosystem thresholds driven by aridity. Science 367, 787–790. 10.1126/science.aay5958

Butinar, L., Spencer-Martins, I., Gunde-Cimerman, N., 2007. Yeasts in high Arctic glaciers: the discovery of a new habitat for eukaryotic microorganisms. Antonie van Leeuwenhoek 91, 277–289. 10.1007/s10482-006-9117-3

Buzzini, P., Turk, M., Perini, L., Turchetti, B., Gunde-Cimerman, N., 2017. Yeasts in Polar and Subpolar Habitats, in: Buzzini, P., Lachance, M.-A., Yurkov, A. (Eds.), Yeasts in Natural Ecosystems: Diversity. Springer International Publishing, Cham, pp. 331–365. 10.1007/978-3-319-62683-3_11

Callahan, B.J., McMurdie, P.J., Rosen, M.J., Han, A.W., Johnson, A.J.A., Holmes, S.P., 2016. DADA2: High-resolution sample inference from Illumina amplicon data. Nat Methods 13, 581–583. 10.1038/nmeth.3869

Cantrell, S.A., Dianese, J.C., Fell, J., Gunde-Cimerman, N., Zalar, P., 2011. Unusual fungal niches. Mycologia 103, 1161–1174. 10.3852/11-108

Carr, E.C., n.d. Characterization of a novel polyextremotolerant fungus, Exophiala viscosa, with insights into its melanin regulation and ecological niche.

Castañeda-Tamez, P., Chiquete-Félix, N., Uribe-Carvajal, S., Cabrera-Orefice, A., 2024. The mitochondrial respiratory chain from Rhodotorula mucilaginosa, an extremophile yeast. Biochimica et Biophysica Acta (BBA) - Bioenergetics 1865, 149035. 10.1016/j.bbabio.2024.149035

Chávez-González, J.D., Flores-Núñez, V.M., Merino-Espinoza, I.U., Partida-Martínez, L.P., 2024. Desert plants, arbuscular mycorrhizal fungi and associated bacteria: Exploring the diversity and role of symbiosis under drought. Environ Microbiol Rep 16, e13300. 10.1111/1758-2229.13300

Chen, K., Liao, H., Arnold, A.E., Korotkin, H.B., Wu, S.H., Matheny, P.B., Lutzoni, F., 2022. Comparative transcriptomics of fungal endophytes in co-culture with their moss host *Dicranum scoparium* reveals fungal trophic lability and moss unchanged to slightly increased growth rates. New Phytologist 234, 1832–1847. 10.1111/nph.18078

Coleine, C., Delgado-Baquerizo, M., DiRuggiero, J., Guirado, E., Harfouche, A.L., Perez-Fernandez, C., Singh, B.K., Selbmann, L., Egidi, E., 2024a. Dryland microbiomes reveal community adaptations to desertification and climate change. The ISME Journal 18, wrae056. 10.1093/ismejo/wrae056

Coleine, C., Kurbessoian, T., Calia, G., Delgado-Baquerizo, M., Cestaro, A., Pindo, M., Armanini, F., Asnicar, F., Isola, D., Segata, N., Donati, C., Stajich, J.E., De Hoog, S., Selbmann, L., 2024b. Class-wide genomic tendency throughout specific extremes in black fungi. Fungal Diversity 125, 121–138. 10.1007/s13225-024-00533-y

Coleine, C., Stajich, J.E., Selbmann, L., 2022. Fungi are key players in extreme ecosystems. Trends in Ecology & Evolution 37, 517–528. 10.1016/j.tree.2022.02.002

Cook, B.I., Ault, T.R., Smerdon, J.E., 2015. Unprecedented 21st century drought risk in the American Southwest and Central Plains. Sci. Adv. 1, e1400082. 10.1126/sciadv.1400082

Dadachova, E., Bryan, R.A., Huang, X., Moadel, T., Schweitzer, A.D., Aisen, P., Nosanchuk, J.D., Casadevall, A., 2007. Ionizing Radiation Changes the Electronic Properties of Melanin and Enhances the Growth of Melanized Fungi. PLoS ONE 2, e457. 10.1371/journal.pone.0000457

Duarte, A.W.F., Dayo-Owoyemi, I., Nobre, F.S., Pagnocca, F.C., Chaud, L.C.S., Pessoa, A., Felipe, M.G.A., Sette, L.D., 2013. Taxonomic assessment and enzymes production by yeasts isolated from marine and terrestrial Antarctic samples. Extremophiles 17, 1023–1035. 10.1007/s00792-013-0584-y

Egidi, E., Delgado-Baquerizo, M., Berdugo, M., Guirado, E., Albanese, D., Singh, B.K., Coleine, C., 2023. UV index and climate seasonality explain fungal community turnover in global drylands. Global Ecol Biogeogr 32, 132–144. 10.1111/geb.13607

Elbert, W., Weber, B., Burrows, S., Steinkamp, J., Büdel, B., Andreae, M.O., Pöschl, U., 2012. Contribution of cryptogamic covers to the global cycles of carbon and nitrogen. Nature Geosci 5, 459–462. 10.1038/ngeo1486

Fenchel, T., 2008. The microbial loop – 25 years later. Journal of Experimental Marine Biology and Ecology 366, 99–103. 10.1016/j.jembe.2008.07.013

Field, K.J., Pressel, S., Duckett, J.G., Rimington, W.R., Bidartondo, M.I., 2015. Symbiotic options for the conquest of land. Trends in Ecology & Evolution 30, 477–486. 10.1016/j.tree.2015.05.007

García-Carmona, M., Lepinay, C., García-Orenes, F., Baldrian, P., Arcenegui, V., Cajthaml, T., Mataix-Solera, J., 2022. Moss biocrust accelerates the recovery and resilience of soil microbial communities in fire-affected semi-arid Mediterranean soils. Science of The Total Environment 846, 157467. 10.1016/j.scitotenv.2022.157467

Gundlapally, S.R., Garcia-Pichel, F., 2006. The Community and Phylogenetic Diversity of Biological Soil Crusts in the Colorado Plateau Studied by Molecular Fingerprinting and Intensive Cultivation. Microb Ecol 52, 345–357. 10.1007/s00248-006-9011-6

Hanke, S.T., Rensing, S.A., n.d. In vitro association of non-seed plant gametophytes with arbuscular mycorrhiza fungi.

Hättenschwiler, S., Coq, S., Barantal, S., Handa, I.T., 2011. Leaf traits and decomposition in tropical rainforests: revisiting some commonly held views and towards a new hypothesis. New Phytologist 189, 950–965. 10.1111/j.1469-8137.2010.03483.x

Hu, W., n.d. Aridity-driven shift in biodiversity–soil multifunctionality relationships.

Huang, J., Li, Y., Fu, C., Chen, F., Fu, Q., Dai, A., Shinoda, M., Ma, Z., Guo, W., Li, Z., Zhang, L., Liu, Y., Yu, H., He, Y., Xie, Y., Guan, X., Ji, M., Lin, L., Wang, S., Yan, H., Wang, G., 2017. Dryland climate change: Recent progress and challenges. Reviews of Geophysics 55, 719–778. 10.1002/2016RG000550

Kapoor, R., Giri, B., Mukerji, K.G., n.d. Glomus macrocarpum: a potential bioinoculant to improve essential oil quality and concentration in Dill (Anethum graveolens L.) and Carum (Trachyspermum ammi (Linn.) Sprague).

Katoh, K., Standley, D.M., 2013. MAFFT Multiple Sequence Alignment Software Version 7: Improvements in Performance and Usability. Molecular Biology and Evolution 30, 772–780. 10.1093/molbev/mst010

Kelly, K.H., Coleine, C., Coshland, C., Stajich, J.E., 2024. Novel Glomeromycotina-Moss Associations Identified in California Dryland Biocrusts. bioRxiv. 10.1101/2024.12.27.630416

Khunnamwong, P., Jindamorakot, S., Limtong, S., 2018. Endophytic yeast diversity in leaf tissue of rice, corn and sugarcane cultivated in Thailand assessed by a culture-dependent approach. Fungal Biology 122, 785–799. 10.1016/j.funbio.2018.04.006

Kurbessoian, T., Ahmed, S.A., Quan, Y., De Hoog, S., Stajich, J.E., 2024. Description of new micro-colonial fungi species *Neophaeococcomyces mojavensis*, Coniosporium tulheliwenetii, and Taxawa tesnikishii cultured from biological soil crusts. 10.1101/2024.06.12.598762

Kurbessoian, T., Pombubpa, N., Coleine, C., Selbmann, L., Pietrasiak, N., Stajich, J.E., 2020. Black Yeast As Desert Sunscreen: Assessing the Genetic Composition of Black Yeasts Found within Biological Soil Crusts.

Ladrón De Guevara, M., Maestre, F.T., 2022. Ecology and responses to climate change of biocrust-forming mosses in drylands. Journal of Experimental Botany 73, 4380–4395. 10.1093/jxb/erac183

León-Sánchez, L., Nicolás, E., Goberna, M., Prieto, I., Maestre, F.T., Querejeta, J.I., 2018. Poor plant performance under simulated climate change is linked to mycorrhizal responses in a semi-arid shrubland. Journal of Ecology 106, 960–976. 10.1111/1365-2745.12888

Li, X., Bai, W., Yang, Q., Yin, B., Zhang, Z., Zhao, B., Kuang, T., Zhang, Y., Zhang, D., 2024. The extremotolerant desert moss Syntrichia caninervis is a promising pioneer plant for colonizing extraterrestrial environments. The Innovation 5, 100657. 10.1016/j.xinn.2024.100657

Love, M.I., Huber, W., Anders, S., 2014. Moderated estimation of fold change and dispersion for RNA-seq data with DESeq2. Genome Biol 15, 550. 10.1186/s13059-014-0550-8

Maestre, F.T., Benito, B.M., Berdugo, M., Concostrina-Zubiri, L., Delgado-Baquerizo, M., Eldridge, D.J., Guirado, E., Gross, N., Kéfi, S., Le Bagousse-Pinguet, Y., Ochoa-Hueso, R., Soliveres, S., 2021. Biogeography of global drylands. New Phytologist 231, 540–558. 10.1111/nph.17395

Maestre, F.T., Delgado-Baquerizo, M., Jeffries, T.C., Eldridge, D.J., Ochoa, V., Gozalo, B., Quero, J.L., García-Gómez, M., Gallardo, A., Ulrich, W., Bowker, M.A., Arredondo, T., Barraza-Zepeda, C., Bran, D., Florentino, A., Gaitán, J., Gutiérrez, J.R., Huber-Sannwald, E., Jankju, M., Mau, R.L., Miriti, M., Naseri, K., Ospina, A., Stavi, I., Wang, D., Woods, N.N., Yuan, X., Zaady, E., Singh, B.K., 2015. Increasing aridity reduces soil microbial diversity and abundance in global drylands. Proc. Natl. Acad. Sci. U.S.A. 112, 15684–15689. 10.1073/pnas.1516684112

Mathieu, D., Bryson, A.E., Hamberger, Britta, Singan, V., Keymanesh, K., Wang, M., Barry, K., Mondo, S., Pangilinan, J., Koriabine, M., Grigoriev, I.V., Bonito, G., Hamberger, Björn, 2024. Multilevel analysis between *Physcomitrium patens* and Mortierellaceae endophytes explores potential long-standing interaction among land plants and fungi. The Plant Journal 118, 304–323. 10.1111/tpj.16605

McMurdie, P.J., Holmes, S., 2014. Waste Not, Want Not: Why Rarefying Microbiome Data Is Inadmissible. PLoS Comput Biol 10, e1003531. 10.1371/journal.pcbi.1003531

McMurdie, P.J., Holmes, S., 2013. phyloseq: An R Package for Reproducible Interactive Analysis and Graphics of Microbiome Census Data. PLoS ONE 8, e61217. 10.1371/journal.pone.0061217

Ochoa-Hueso, R., Eldridge, D.J., Delgado-Baquerizo, M., Soliveres, S., Bowker, M.A., Gross, N., Le Bagousse-Pinguet, Y., Quero, J.L., García-Gómez, M., Valencia, E., Arredondo, T., Beinticinco, L., Bran, D., Cea, A., Coaguila, D., Dougill, A.J., Espinosa, C.I., Gaitán, J., Guuroh, R.T., Guzman, E., Gutiérrez, J.R., Hernández, R.M., Huber-Sannwald, E., Jeffries, T., Linstädter, A., Mau, R.L., Monerris, J., Prina, A., Pucheta, E., Stavi, I., Thomas, A.D., Zaady, E., Singh, B.K., Maestre, F.T., 2018. Soil fungal abundance and plant functional traits drive fertile island formation in global drylands. Journal of Ecology 106, 242–253. 10.1111/1365-2745.12871

Oksanen, J., Simpson, G.L., Blanchet, F.G., Kindt, R., Legendre, P., Minchin, P.R., O’Hara, R.B., Solymos, P., Stevens, M.H.H., Szoecs, E., Wagner, H., Barbour, M., Bedward, M., Bolker, B., Borcard, D., Carvalho, G., Chirico, M., De Caceres, M., Durand, S., Evangelista, H.B.A., FitzJohn, R., Friendly, M., Furneaux, B., Hannigan, G., Hill, M.O., Lahti, L., McGlinn, D., Ouellette, M.-H., Ribeiro Cunha, E., Smith, T., Stier, A., Ter Braak, C.J.F., Weedon, J., 2001. vegan: Community Ecology Package. 10.32614/CRAN.package.vegan

Palmer, J.M., Jusino, M.A., Banik, M.T., Lindner, D.L., 2018. Non-biological synthetic spike-in controls and the AMPtk software pipeline improve mycobiome data. PeerJ 6, e4925. 10.7717/peerj.4925

Peng, X., Wang, Y., Tang, L.J., Li, X.X., Xiao, Y.W., Zhang, Z.B., Yan, R.M., Yang, H.L., Chang, J., Zhu, B., Zhu, D., 2018. Yeasts from Nanfeng mandarin plants: occurrence, diversity and capability to produce indole-3-acetic acid. Biotechnology & Biotechnological Equipment 32, 1496–1506. 10.1080/13102818.2018.1487337

Phillips, M.L., McNellis, B.E., Howell, A., Lauria, C.M., Belnap, J., Reed, S.C., 2022. Biocrusts mediate a new mechanism for land degradation under a changing climate. Nat. Clim. Chang. 12, 71–76. 10.1038/s41558-021-01249-6

Pombubpa, N., Pietrasiak, N., De Ley, P., Stajich, J.E., 2020. Insights into dryland biocrust microbiome: geography, soil depth and crust type affect biocrust microbial communities and networks in Mojave Desert, USA. FEMS Microbiology Ecology 96, fiaa125. 10.1093/femsec/fiaa125

Price, M.N., Dehal, P.S., Arkin, A.P., 2010. FastTree 2 – Approximately Maximum-Likelihood Trees for Large Alignments. PLoS ONE 5, e9490. 10.1371/journal.pone.0009490

Pulschen, A.A., Rodrigues, F., Duarte, R.T.D., Araujo, G.G., Santiago, I.F., Paulino-Lima, I.G., Rosa, C.A., Kato, M.J., Pellizari, V.H., Galante, D., 2015. UV-resistant yeasts isolated from a high-altitude volcanic area on the Atacama Desert as eukaryotic models for astrobiology. MicrobiologyOpen 4, 574–588. 10.1002/mbo3.262

Ranzoni, F.V., n.d. Fungi Isolated In Culture From Soils of the Sonoran Desert.

Remke, M.J., Johnson, N.C., Wright, J., Williamson, M., Bowker, M.A., 2021. Sympatric pairings of dryland grass populations, mycorrhizal fungi and associated soil biota enhance mutualism and ameliorate drought stress. Journal of Ecology 109, 1210–1223. 10.1111/1365-2745.13546

Rodriguez-Caballero, E., Belnap, J., Büdel, B., Crutzen, P.J., Andreae, M.O., Pöschl, U., Weber, B., 2018. Dryland photoautotrophic soil surface communities endangered by global change. Nature Geosci 11, 185–189. 10.1038/s41561-018-0072-1

Rodriguez-Caballero, E., Stanelle, T., Egerer, S., Cheng, Y., Su, H., Canton, Y., Belnap, J., Andreae, M.O., Tegen, I., Reick, C.H., Pöschl, U., Weber, B., 2022. Global cycling and climate effects of aeolian dust controlled by biological soil crusts. Nat. Geosci. 15, 458–463. 10.1038/s41561-022-00942-1

Rognes, T., Flouri, T., Nichols, B., Quince, C., Mahé, F., 2016. VSEARCH: a versatile open source tool for metagenomics. PeerJ 4, e2584. 10.7717/peerj.2584

Rudgers, J.A., Dettweiler-Robinson, E., Belnap, J., Green, L.E., Sinsabaugh, R.L., Young, K.E., Cort, C.E., Darrouzet-Nardi, A., 2018. Are fungal networks key to dryland primary production? American J of Botany 105, 1783–1787. 10.1002/ajb2.1184

Russel88/MicEco: v0.9.15 [WWW Document], n.d. URL https://zenodo.org/records/4733747 (accessed 5.30.24).

Schmidt, S.K., Vimercati, L., Darcy, J.L., Arán, P., Gendron, E.M.S., Solon, A.J., Porazinska, D., Dorador, C., 2017. A *Naganishia* in high places: functioning populations or dormant cells from the atmosphere? Mycology 8, 153–163. 10.1080/21501203.2017.1344154

Schoch, C.L., Seifert, K.A., Huhndorf, S., Robert, V., Spouge, J.L., Levesque, C.A., Chen, W., Fungal Barcoding Consortium, Fungal Barcoding Consortium Author List, Bolchacova, E., Voigt, K., Crous, P.W., Miller, A.N., Wingfield, M.J., Aime, M.C., An, K.-D., Bai, F.-Y., Barreto, R.W., Begerow, D., Bergeron, M.-J., Blackwell, M., Boekhout, T., Bogale, M., Boonyuen, N., Burgaz, A.R., Buyck, B., Cai, L., Cai, Q., Cardinali, G., Chaverri, P., Coppins, B.J., Crespo, A., Cubas, P., Cummings, C., Damm, U., De Beer, Z.W., De Hoog, G.S., Del-Prado, R., Dentinger, B., Diéguez-Uribeondo, J., Divakar, P.K., Douglas, B., Dueñas, M., Duong, T.A., Eberhardt, U., Edwards, J.E., Elshahed, M.S., Fliegerova, K., Furtado, M., García, M.A., Ge, Z.-W., Griffith, G.W., Griffiths, K., Groenewald, J.Z., Groenewald, M., Grube, M., Gryzenhout, M., Guo, L.-D., Hagen, F., Hambleton, S., Hamelin, R.C., Hansen, K., Harrold, P., Heller, G., Herrera, C., Hirayama, K., Hirooka, Y., Ho, H.-M., Hoffmann, K., Hofstetter, V., Högnabba, F., Hollingsworth, P.M., Hong, S.-B., Hosaka, K., Houbraken, J., Hughes, K., Huhtinen, S., Hyde, K.D., James, T., Johnson, E.M., Johnson, J.E., Johnston, P.R., Jones, E.B.G., Kelly, L.J., Kirk, P.M., Knapp, D.G., Kõljalg, U., Kovács, G.M., Kurtzman, C.P., Landvik, S., Leavitt, S.D., Liggenstoffer, A.S., Liimatainen, K., Lombard, L., Luangsa-ard, J.J., Lumbsch, H.T., Maganti, H., Maharachchikumbura, S.S.N., Martin, M.P., May, T.W., McTaggart, A.R., Methven, A.S., Meyer, W., Moncalvo, J.-M., Mongkolsamrit, S., Nagy, L.G., Nilsson, R.H., Niskanen, T., Nyilasi, I., Okada, G., Okane, I., Olariaga, I., Otte, J., Papp, T., Park, D., Petkovits, T., Pino-Bodas, R., Quaedvlieg, W., Raja, H.A., Redecker, D., Rintoul, T.L., Ruibal, C., Sarmiento-Ramírez, J.M., Schmitt, I., Schüßler, A., Shearer, C., Sotome, K., Stefani, F.O.P., Stenroos, S., Stielow, B., Stockinger, H., Suetrong, S., Suh, S.-O., Sung, G.-H., Suzuki, M., Tanaka, K., Tedersoo, L., Telleria, M.T., Tretter, E., Untereiner, W.A., Urbina, H., Vágvölgyi, C., Vialle, A., Vu, T.D., Walther, G., Wang, Q.-M., Wang, Y., Weir, B.S., Weiß, M., White, M.M., Xu, J., Yahr, R., Yang, Z.L., Yurkov, A., Zamora, J.-C., Zhang, N., Zhuang, W.-Y., Schindel, D., 2012. Nuclear ribosomal internal transcribed spacer (ITS) region as a universal DNA barcode marker for *Fungi*. Proc. Natl. Acad. Sci. U.S.A. 109, 6241–6246. 10.1073/pnas.1117018109

Segal-Kischinevzky, C., Romero-Aguilar, L., Alcaraz, L.D., López-Ortiz, G., Martínez-Castillo, B., Torres-Ramírez, N., Sandoval, G., González, J., 2022. Yeasts Inhabiting Extreme Environments and Their Biotechnological Applications. Microorganisms 10, 794. 10.3390/microorganisms10040794

Selbmann, L., Benkő, Z., Coleine, C., De Hoog, S., Donati, C., Druzhinina, I., Emri, T., Ettinger, C.L., Gladfelter, A.S., Gorbushina, A.A., Grigoriev, I.V., Grube, M., Gunde-Cimerman, N., Karányi, Z.Á., Kocsis, B., Kubressoian, T., Miklós, I., Miskei, M., Muggia, L., Northen, T., Novak-Babič, M., Pennacchio, C., Pfliegler, W.P., Pòcsi, I., Prigione, V., Riquelme, M., Segata, N., Schumacher, J., Shelest, E., Sterflinger, K., Tesei, D., U’Ren, J.M., Varese, G.C., Vázquez-Campos, X., Vicente, V.A., Souza, E.M., Zalar, P., Walker, A.K., Stajich, J.E., 2020. Shed Light in the DaRk LineagES of the Fungal Tree of Life—STRES. Life 10, 362. 10.3390/life10120362

Selbmann, L., de Hoog, G.S., Mazzaglia, A., Friedmann, E.I., Onofri, S., n.d. Fungi at the edge of life: cryptoendolithic black fungi from Antarctic desert.

Selbmann, L., Isola, D., Zucconi, L., Onofri, S., 2011. Resistance to UV-B induced DNA damage in extreme-tolerant cryptoendolithic Antarctic fungi: detection by PCR assays. Fungal Biology 115, 937–944. 10.1016/j.funbio.2011.02.016

Selbmann, L., Stoppiello, G.A., Onofri, S., Stajich, J.E., Coleine, C., 2021. Culture-Dependent and Amplicon Sequencing Approaches Reveal Diversity and Distribution of Black Fungi in Antarctic Cryptoendolithic Communities. JoF 7, 213. 10.3390/jof7030213

Selbmann, L., Zucconi, L., Onofri, S., Cecchini, C., Isola, D., Turchetti, B., Buzzini, P., 2014. Taxonomic and phenotypic characterization of yeasts isolated from worldwide cold rock-associated habitats. Fungal Biology 118, 61–71. 10.1016/j.funbio.2013.11.002

Sen, D., Paul, K., Saha, C., Mukherjee, G., Nag, M., Ghosh, S., Das, A., Seal, A., Tripathy, S., 2019. A unique life-strategy of an endophytic yeast Rhodotorula mucilaginosa JGTA-S1—a comparative genomics viewpoint. DNA Research 26, 131–146. 10.1093/dnares/dsy044

Silambarasan, S., Logeswari, P., Cornejo, P., Kannan, V.R., 2019. Evaluation of the production of exopolysaccharide by plant growth promoting yeast Rhodotorula sp. strain CAH2 under abiotic stress conditions. International Journal of Biological Macromolecules 121, 55–62. 10.1016/j.ijbiomac.2018.10.016

Silva, A.T., Gao, B., Fisher, K.M., Mishler, B.D., Ekwealor, J.T.B., Stark, L.R., Li, X., Zhang, D., Bowker, M.A., Brinda, J.C., Coe, K.K., Oliver, M.J., 2021. To dry perchance to live: Insights from the genome of the desiccation-tolerant biocrust moss *Syntrichia caninervis*. The Plant Journal 105, 1339–1356. 10.1111/tpj.15116

Smith, M.E., Henkel, T.W., Williams, G.C., Aime, M.C., Fremier, A.K., Vilgalys, R., 2017. Investigating niche partitioning of ectomycorrhizal fungi in specialized rooting zones of the monodominant leguminous tree *Dicymbe corymbosa*. New Phytologist 215, 443–453. 10.1111/nph.14570

Sterflinger, K., Tesei, D., Zakharova, K., 2012. Fungi in hot and cold deserts with particular reference to microcolonial fungi. Fungal Ecology 5, 453–462. 10.1016/j.funeco.2011.12.007

Sun, X., Sharon, O., Sharon, A., 2023. Distinct Features Based on Partitioning of the Endophytic Fungi of Cereals and Other Grasses. Microbiol Spectr 11, e00611–23. 10.1128/spectrum.00611-23

Talbot, J.M., Bruns, T.D., Taylor, J.W., Smith, D.P., Branco, S., Glassman, S.I., Erlandson, S., Vilgalys, R., Liao, H.-L., Smith, M.E., Peay, K.G., 2014. Endemism and functional convergence across the North American soil mycobiome. Proc. Natl. Acad. Sci. U.S.A. 111, 6341–6346. 10.1073/pnas.1402584111

Taudière, A., 2023. MiscMetabar: an R package to facilitate visualizationand reproducibility in metabarcoding analysis. JOSS 8, 6038. 10.21105/joss.06038

Thompson, L.R., Sanders, J.G., McDonald, D., Amir, A., Ladau, J., Locey, K.J., Prill, R.J., Tripathi, A., Gibbons, S.M., Ackermann, G., Navas-Molina, J.A., Janssen, S., Kopylova, E., Vázquez-Baeza, Y., González, A., Morton, J.T., Mirarab, S., Zech Xu, Z., Jiang, L., Haroon, M.F., Kanbar, J., Zhu, Q., Jin Song, S., Kosciolek, T., Bokulich, N.A., Lefler, J., Brislawn, C.J., Humphrey, G., Owens, S.M., Hampton-Marcell, J., Berg-Lyons, D., McKenzie, V., Fierer, N., Fuhrman, J.A., Clauset, A., Stevens, R.L., Shade, A., Pollard, K.S., Goodwin, K.D., Jansson, J.K., Gilbert, J.A., Knight, R., The Earth Microbiome Project Consortium, Rivera, J.L.A., Al-Moosawi, L., Alverdy, J., Amato, K.R., Andras, J., Angenent, L.T., Antonopoulos, D.A., Apprill, A., Armitage, D., Ballantine, K., Bárta, J., Baum, J.K., Berry, A., Bhatnagar, A., Bhatnagar, M., Biddle, J.F., Bittner, L., Boldgiv, B., Bottos, E., Boyer, D.M., Braun, J., Brazelton, W., Brearley, F.Q., Campbell, A.H., Caporaso, J.G., Cardona, C., Carroll, J., Cary, S.C., Casper, B.B., Charles, T.C., Chu, H., Claar, D.C., Clark, R.G., Clayton, J.B., Clemente, J.C., Cochran, A., Coleman, M.L., Collins, G., Colwell, R.R., Contreras, M., Crary, B.B., Creer, S., Cristol, D.A., Crump, B.C., Cui, D., Daly, S.E., Davalos, L., Dawson, R.D., Defazio, J., Delsuc, F., Dionisi, H.M., Dominguez-Bello, M.G., Dowell, R., Dubinsky, E.A., Dunn, P.O., Ercolini, D., Espinoza, R.E., Ezenwa, V., Fenner, N., Findlay, H.S., Fleming, I.D., Fogliano, V., Forsman, A., Freeman, C., Friedman, E.S., Galindo, G., Garcia, L., Garcia-Amado, M.A., Garshelis, D., Gasser, R.B., Gerdts, G., Gibson, M.K., Gifford, I., Gill, R.T., Giray, T., Gittel, A., Golyshin, P., Gong, D., Grossart, H.-P., Guyton, K., Haig, S.-J., Hale, V., Hall, R.S., Hallam, S.J., Handley, K.M., Hasan, N.A., Haydon, S.R., Hickman, J.E., Hidalgo, G., Hofmockel, K.S., Hooker, J., Hulth, S., Hultman, J., Hyde, E., Ibáñez-Álamo, J.D., Jastrow, J.D., Jex, A.R., Johnson, L.S., Johnston, E.R., Joseph, S., Jurburg, S.D., Jurelevicius, D., Karlsson, A., Karlsson, R., Kauppinen, S., Kellogg, C.T.E., Kennedy, S.J., Kerkhof, L.J., King, G.M., Kling, G.W., Koehler, A.V., Krezalek, M., Kueneman, J., Lamendella, R., Landon, E.M., Lane-deGraaf, K., LaRoche, J., Larsen, P., Laverock, B., Lax, S., Lentino, M., Levin, I.I., Liancourt, P., Liang, W., Linz, A.M., Lipson, D.A., Liu, Y., Lladser, M.E., Lozada, M., Spirito, C.M., MacCormack, W.P., MacRae-Crerar, A., Magris, M., Martín-Platero, A.M., Martín-Vivaldi, M., Martínez, L.M., Martínez-Bueno, M., Marzinelli, E.M., Mason, O.U., Mayer, G.D., McDevitt-Irwin, J.M., McDonald, J.E., McGuire, K.L., McMahon, K.D., McMinds, R., Medina, M., Mendelson, J.R., Metcalf, J.L., Meyer, F., Michelangeli, F., Miller, K., Mills, D.A., Minich, J., Mocali, S., Moitinho-Silva, L., Moore, A., Morgan-Kiss, R.M., Munroe, P., Myrold, D., Neufeld, J.D., Ni, Y., Nicol, G.W., Nielsen, S., Nissimov, J.I., Niu, K., Nolan, M.J., Noyce, K., O’Brien, S.L., Okamoto, N., Orlando, L., Castellano, Y.O., Osuolale, O., Oswald, W., Parnell, J., Peralta-Sánchez, J.M., Petraitis, P., Pfister, C., Pilon-Smits, E., Piombino, P., Pointing, S.B., Pollock, F.J., Potter, C., Prithiviraj, B., Quince, C., Rani, A., Ranjan, R., Rao, S., Rees, A.P., Richardson, M., Riebesell, U., Robinson, C., Rockne, K.J., Rodriguezl, S.M., Rohwer, F., Roundstone, W., Safran, R.J., Sangwan, N., Sanz, V., Schrenk, M., Schrenzel, M.D., Scott, N.M., Seger, R.L., Seguin-Orlando, A., Seldin, L., Seyler, L.M., Shakhsheer, B., Sheets, G.M., Shen, C., Shi, Y., Shin, H., Shogan, B.D., Shutler, D., Siegel, J., Simmons, S., Sjöling, S., Smith, D.P., Soler, J.J., Sperling, M., Steinberg, P.D., Stephens, B., Stevens, M.A., Taghavi, S., Tai, V., Tait, K., Tan, C.L., Taş, N., Taylor, D.L., Thomas, T., Timling, I., Turner, B.L., Urich, T., Ursell, L.K., Van Der Lelie, D., Van Treuren, W., Van Zwieten, L., Vargas-Robles, D., Thurber, R.V., Vitaglione, P., Walker, D.A., Walters, W.A., Wang, S., Wang, T., Weaver, T., Webster, N.S., Wehrle, B., Weisenhorn, P., Weiss, S., Werner, J.J., West, K., Whitehead, A., Whitehead, S.R., Whittingham, L.A., Willerslev, E., Williams, A.E., Wood, S.A., Woodhams, D.C., Yang, Y., Zaneveld, J., Zarraonaindia, I., Zhang, Q., Zhao, H., 2017. A communal catalogue reveals Earth’s multiscale microbial diversity. Nature 551, 457–463. 10.1038/nature24621

Trejos-Espeleta, J.C., Juan, M.-J., Schmidt, S., Summers, P., Bradley, J., Orsi, W., 2024. Principal role of fungi in soil carbon stabilization during early pedogenesis in the high Arctic. Proceedings of the National Academy of Sciences. 10.1073/pnas.2402689121

U’Ren, J.M., Lutzoni, F., Miadlikowska, J., Arnold, A.E., 2010. Community Analysis Reveals Close Affinities Between Endophytic and Endolichenic Fungi in Mosses and Lichens. Microb Ecol 60, 340–353. 10.1007/s00248-010-9698-2

Wang, L.-W., Xu, B.-G., Wang, J.-Y., Su, Z.-Z., Lin, F.-C., Zhang, C.-L., Kubicek, C.P., 2012. Bioactive metabolites from Phoma species, an endophytic fungus from the Chinese medicinal plant Arisaema erubescens. Appl Microbiol Biotechnol 93, 1231–1239. 10.1007/s00253-011-3472-3

Warren, S.D., Clair, L.L., Stark, L.R., Lewis, L.A., Pombubpa, N., Kurbessoian, T., Stajich, J.E., Aanderud, Z.T., 2019. Reproduction and Dispersal of Biological Soil Crust Organisms. Front. Ecol. Evol. 7, 344. 10.3389/fevo.2019.00344

Weber, B., Belnap, J., Büdel, B., Antoninka, A.J., Barger, N.N., Chaudhary, V.B., Darrouzet-Nardi, A., Eldridge, D.J., Faist, A.M., Ferrenberg, S., Havrilla, C.A., Huber-Sannwald, E., Malam Issa, O., Maestre, F.T., Reed, S.C., Rodriguez-Caballero, E., Tucker, C., Young, K.E., Zhang, Y., Zhao, Y., Zhou, X., Bowker, M.A., 2022. What is a biocrust? A refined, contemporary definition for a broadening research community. Biological Reviews 97, 1768–1785. 10.1111/brv.12862

Weber, B., Büdel, B., Belnap, J. (Eds.), 2016. Biological Soil Crusts: An Organizing Principle in Drylands, Ecological Studies. Springer International Publishing, Cham. 10.1007/978-3-319-30214-0

Wei, X.-Y., Zhu, H.-Y., Song, L., Zhang, R.-P., Li, A.-H., Niu, Q.-H., Liu, X.-Z., Bai, F.-Y., 2022.Yeast Diversity in the Qaidam Basin Desert in China with the Description of Five New Yeast Species. JoF 8, 858. 10.3390/jof8080858

Weiss, J.L., Overpeck, J.T., 2005. Is the Sonoran Desert losing its cool? Global Change Biology 11, 2065–2077. 10.1111/j.1365-2486.2005.01020.x

White, Bruns, T., Lee, S., Taylor, J., 1990. White, T. J., T. D. Bruns, S. B. Lee, and J. W. Taylor. Amplification and direct sequencing of fungal ribosomal RNA Genes for phylogenetics. pp. 315– 322.

Xu, S., Li, L., Luo, X., Chen, M., Tang, W., Zhan, L., Dai, Z., Lam, T.T., Guan, Y., Yu, G., 2022. *Ggtree*: A serialized data object for visualization of a phylogenetic tree and annotation data. iMeta 1, e56. 10.1002/imt2.56

Zakharova, K., Tesei, D., Marzban, G., Dijksterhuis, J., Wyatt, T., Sterflinger, K., 2013. Microcolonial Fungi on Rocks: A Life in Constant Drought? Mycopathologia 175, 537–547. 10.1007/s11046-012-9592-1

Zhang, T., Jia, R.-L., Yu, L.-Y., 2016. Diversity and distribution of soil fungal communities associated with biological soil crusts in the southeastern Tengger Desert (China) as revealed by 454 pyrosequencing. Fungal Ecology 23, 156–163. 10.1016/j.funeco.2016.08.004

Zhang, X., Evans, J.P., Burrell, A.L., 2024. Less than 4% of dryland areas are projected to desertify despite increased aridity under climate change. Commun Earth Environ 5, 300. 10.1038/s43247-024-01463-y

