## Supplementary data for "Increased aridity is associated with diversity and composition changes in the biocrust mycobiome"

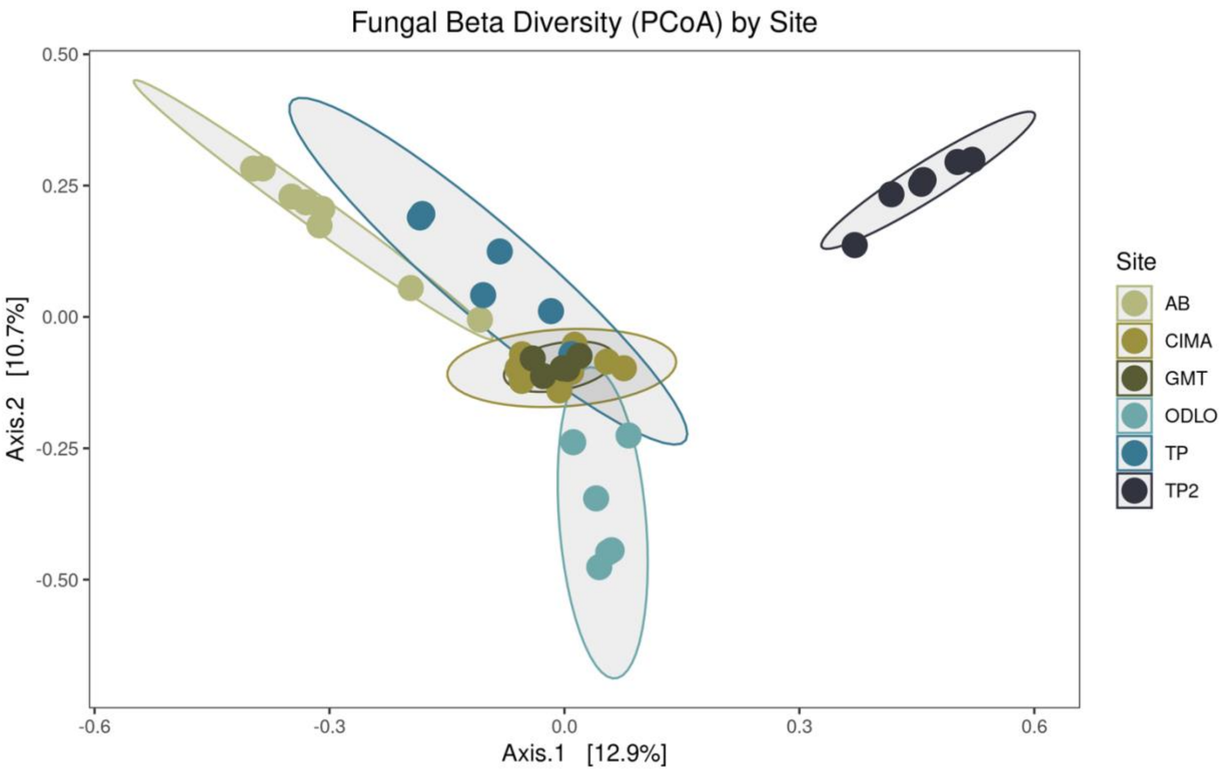


**Figure S1:** Biocrust fungal beta diversity (Bray Curtis Dissimilarity) by site. TP = Torrey Pines, ODLO = Oasis De Los Osos Reserve, AB = Anza Borrego Research Station, CIMA = CIMA Volcanic Field, GMT = Granite Mountains Research Center.

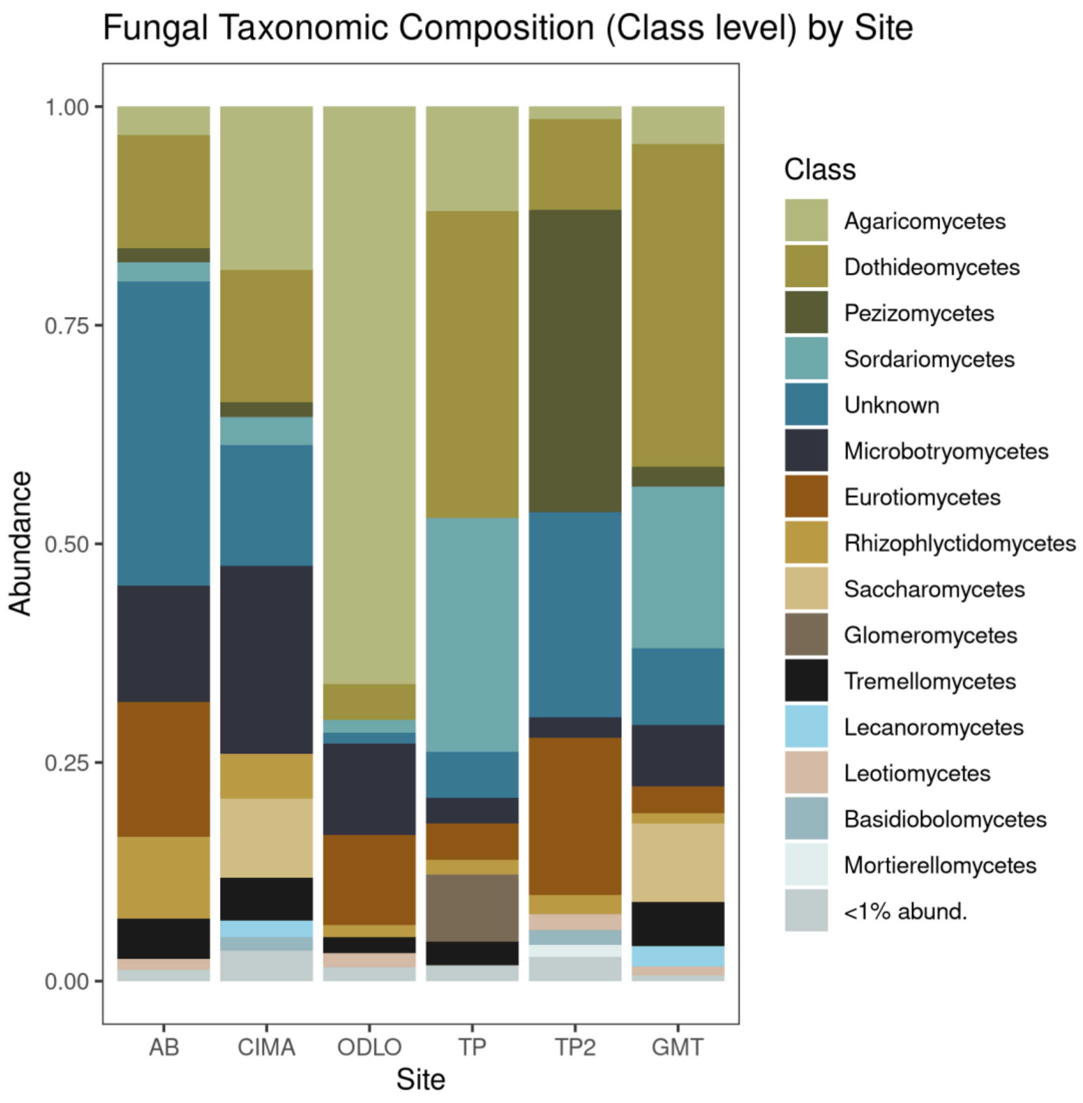


**Figure S2:** Biocrust fungal taxonomic composition by site. TP = Torrey Pines, ODLO = Oasis De Los Osos Reserve, AB = Anza Borrego Research Station, CIMA = CIMA Volcanic Field, GMT = Granite Mountains Research Center.

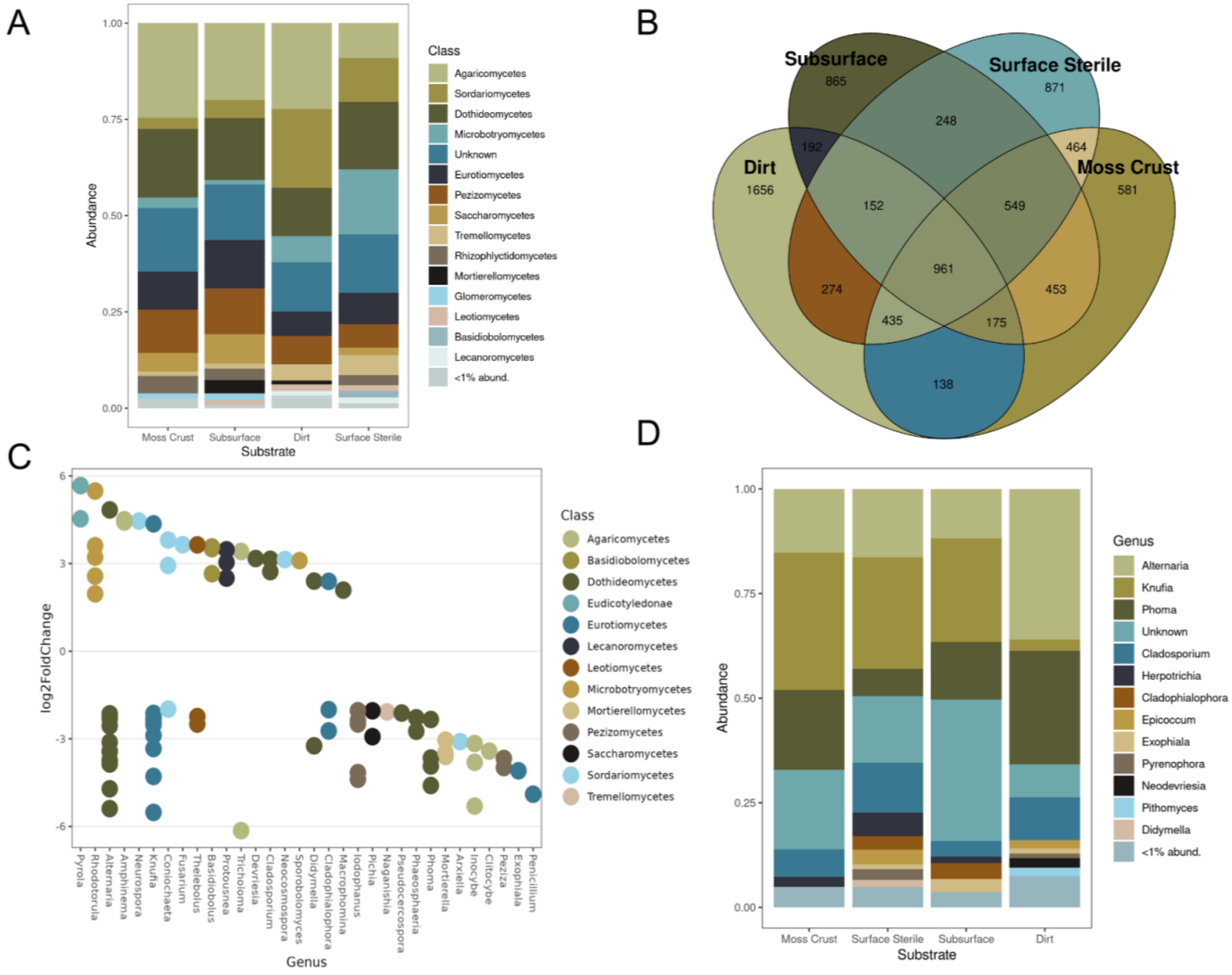


**Figure S3:** (A) Fungal taxonomic composition by substrate. Shifts in relative abundance at the class level are apparent. (B) Venn diagram of ASVs shared by or unique to different substrates. Notably, surface sterile moss has a high level of unique taxa. (C) Significant shifts in relative abundance of different taxa before and after surface sterilization calculated by DEseq. (D) Black yeast taxonomic composition by substrate. Clear differences in taxonomic composition and relative abundance are apparent.

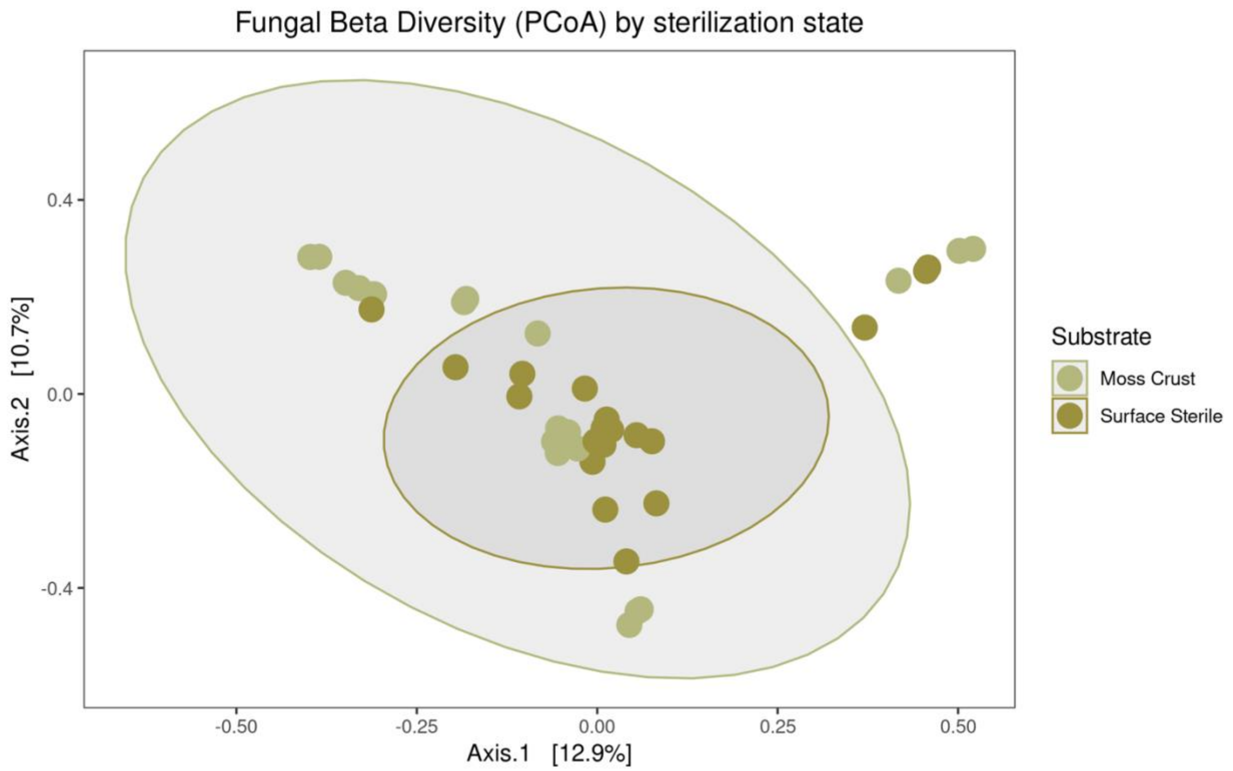


**Figure S4:** Biocrust fungal beta diversity (Bray Curtis Dissimilarity) by sterilization state.

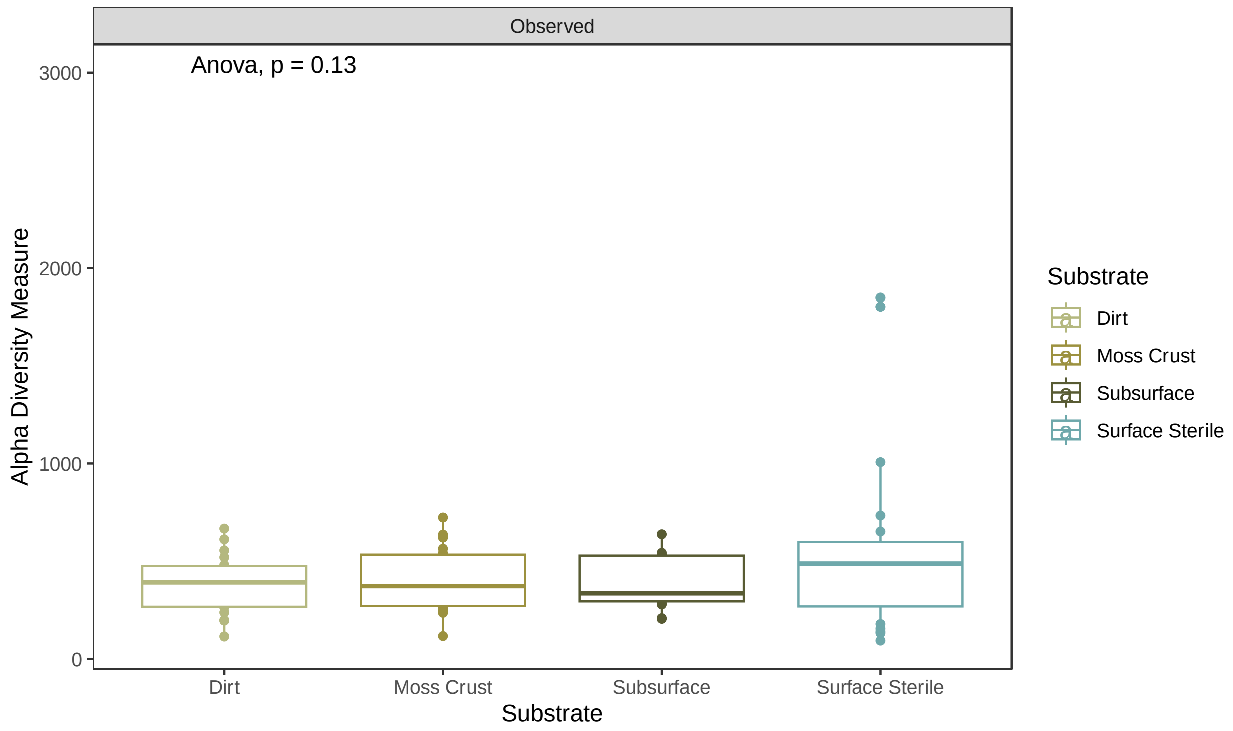


**Figure S5:** Observed alpha diversity by substrate

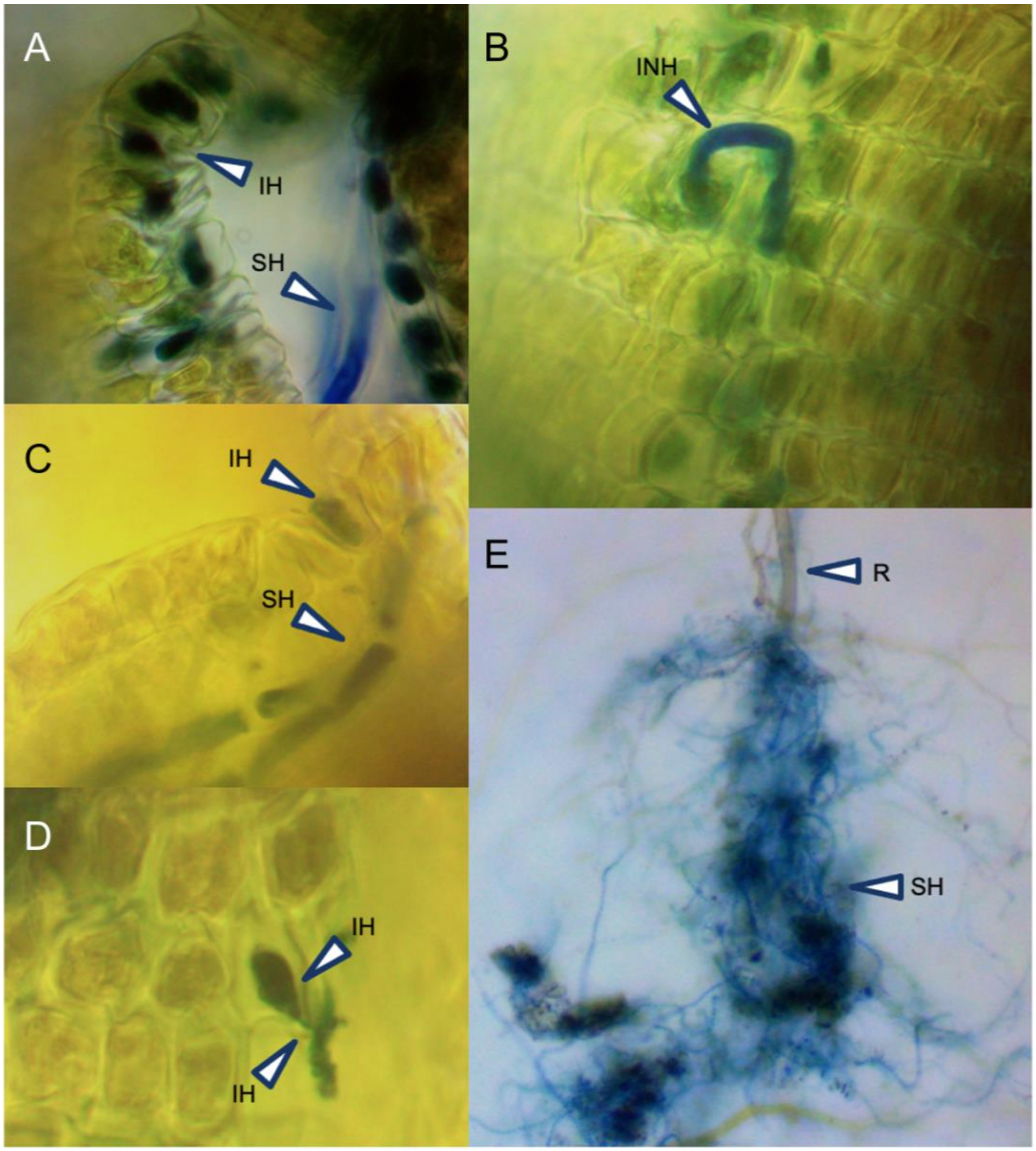


**Figure S6:** Colonization of *Crossidium squamiferum* obtained from Anza Borrego Research Station by dark septate endophytes. (A) *C. squamiferum* gametophyte with healthy photosynthetic tissue (B). Healthy photosynthetic tissue is colonized by dark septate endophytes (C), (D), (E), (F). Septate fungal hyphae wrap around rhizoids within moss biocrusts (G) and move up to colonize the stems and leaves of the moss. IH = Intracellular hyphae, SH = Septate hyphae, INH = Intercellular hyphae, R = Rhizoid.

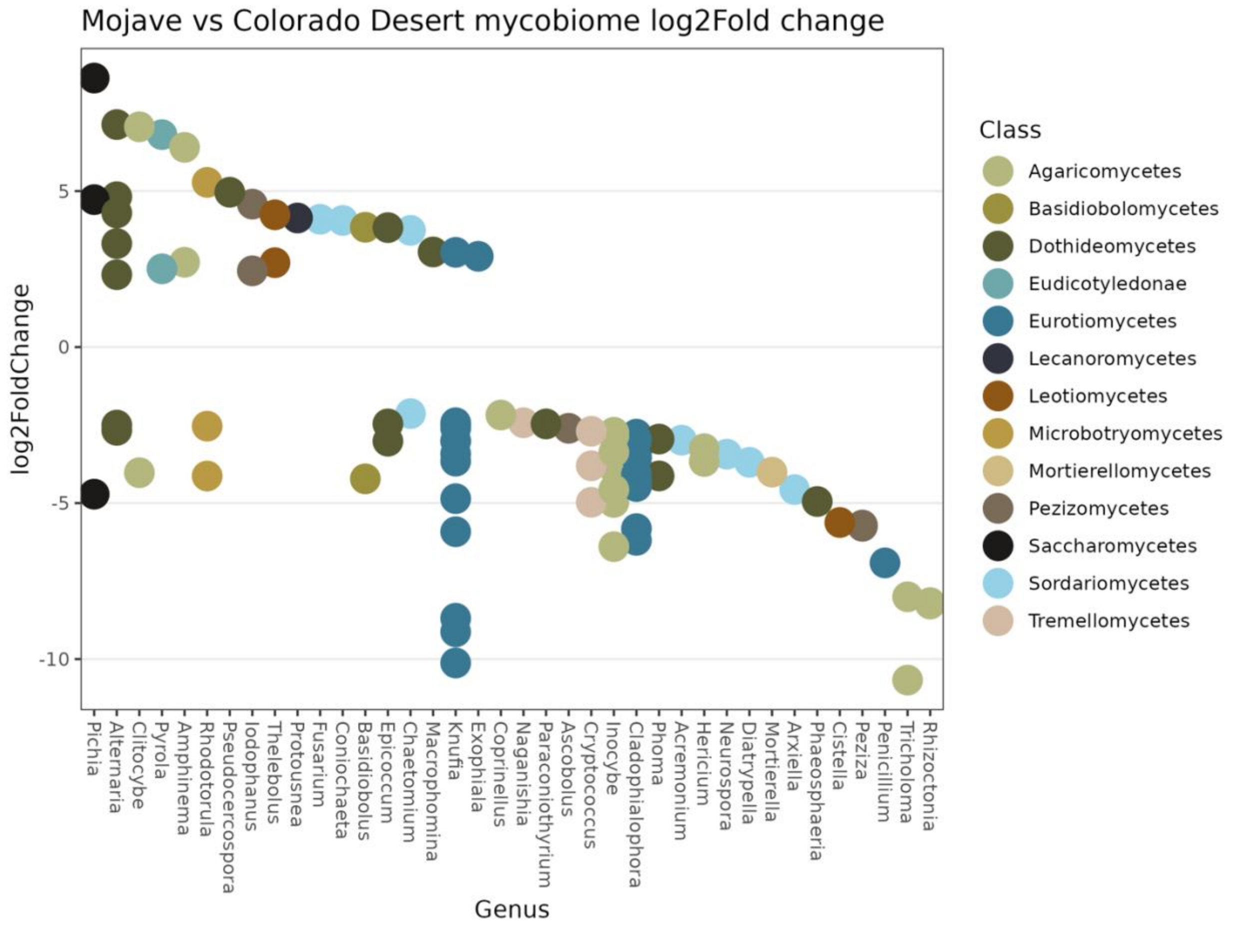


**Figure S7**: Significant relative abundance changes in fungal taxa between the Mojave and Colorado Deserts

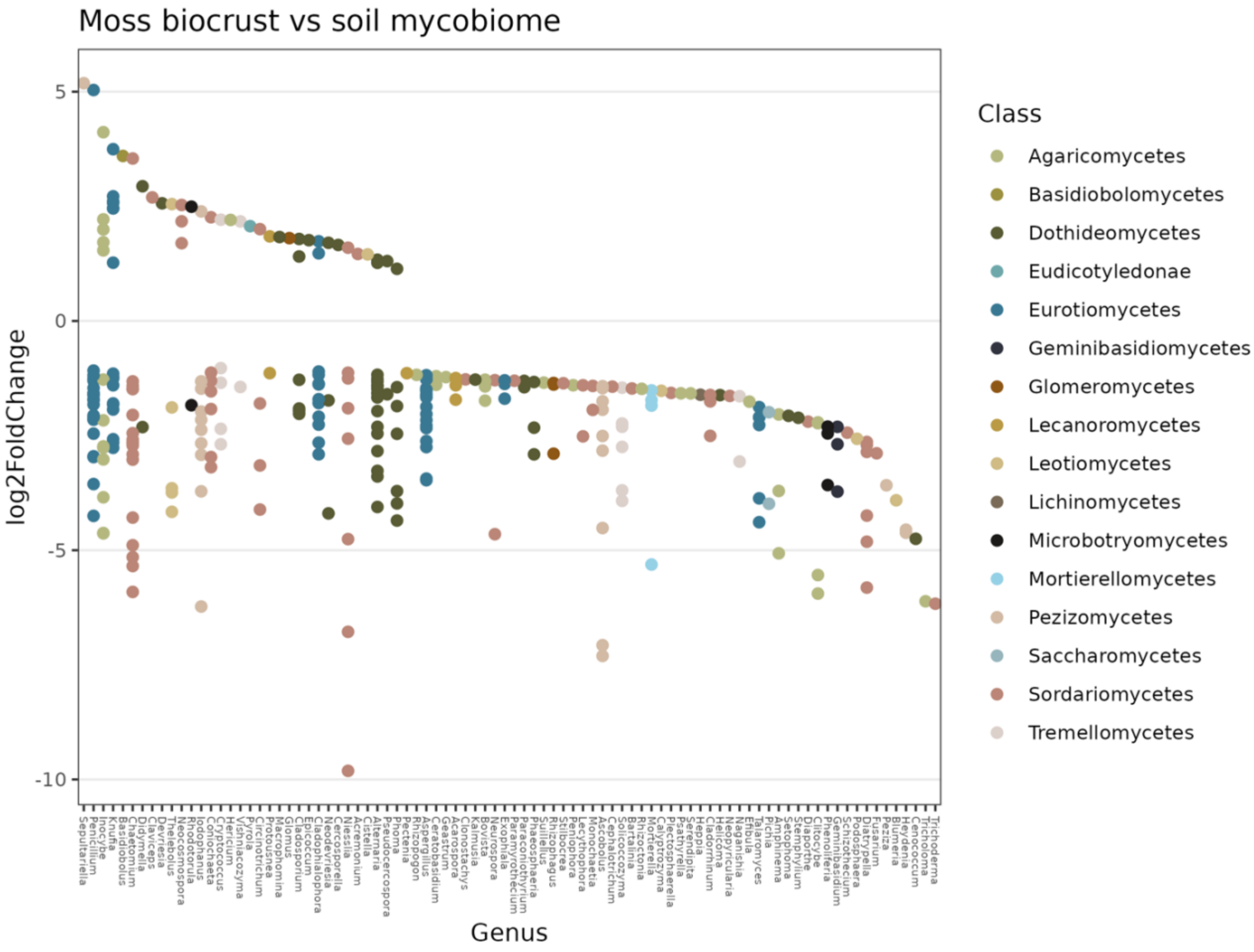


**Figure S8**: Significant relative abundance changes in fungal taxa between the moss biocrust and soil samples.

**Table S1:** Site coordinates and moss taxonomy

| Site | Full site name | Climate | Aridity | Moss family | Moss species | Soil type | Longitude | Latitude |
|---|---|---|---|---|---|---|---|---|
| TP | Torrey Pines Site 1 | Coastal | Semi arid | Pottiaceae | *Trichostomopsis australasiae* | Sandstone | -117.2482757 | 32.9421506 |
| TP2 | Torrey Pines Site 2 | Coastal | Semi arid | Pottiaceae | *Vinealobryum brachyphyllum* | Sandy | -117.2482757 | 32.9421506 |
| AB | Anza Borrego Research Center | Colorado Desert | Arid | Pottiaceae | *Crossidium spp. (squamiferum dominant)* | Sandy | -116.3895333 | 33.2407889 |
| ODLO | Oasis De Los Osos Preserve | Colorado Desert | Arid | Pottiaceae | *Trichostomopsis australasiae* | Granitic | -116.68908 | 33.89182 |
| GMT | Granite Mountains Research Station | Mojave Desert | Hyper arid | Grimiaceae | *Grimia sp.* | Granitic | -115.6297361 | 34.7804861 |
| CIMA | Cima Volcanic Field | Mojave Desert | Hyper arid | Pottiaceae | *Trichostomopsis australasiae* | Volcanic | -115.8699722 | 35.1997917 |

**Table S2:** List of fungal species with differential abundance between semi-arid (coastal) and arid to hyper-arid (Colorado and Mojave Desert) climates. Note, fungi that could not be identified to the species level were filtered from this table.

| ASV | log2FoldChange | pvalue | Phylum | Class | *Species* |
|---|---|---|---|---|---|
| ASV3753 | -6.296 | 5.53E-22 | Ascomycota | Sordariomycetes | *Chaetomium globosum* |
| ASV147 | -4.482 | 2.28E-07 | Ascomycota | Dothideomycetes | *Alternaria tenuissima* |
| ASV1539 | -4.426 | 9.22E-13 | Ascomycota | Eurotiomycetes | *Phaeomoniella chlamydospora* |
| ASV349 | -4.124 | 3.18E-08 | Basidiobolomycota | Basidiobolomycetes | *Basidiobolus ranarum* |
| ASV162 | -3.868 | 3.62E-08 | Ascomycota | Dothideomycetes | *Cercosporella tinosporae* |
| ASV2146 | -3.779 | 5.45E-11 | Ascomycota | Taphrinomycetes | *Taphrina wiesneri* |
| ASV489 | -3.264 | 4.64E-07 | Glomeromycota | Glomeromycetes | *Glomus macrocarpum* |
| ASV7086 | -2.781 | 1.27E-07 | Ascomycota | Eurotiomycetes | *Phaeomoniella chlamydospora* |
| ASV9 | -2.728 | 0.0001362146783 | Ascomycota | Sordariomycetes | *Neocosmospora falciformis* |
| ASV1521 | -2.692 | 1.36E-05 | Ascomycota | Sordariomycetes | *Neocosmospora falciformis* |
| ASV748 | -2.671 | 9.97E-06 | Basidiobolomycota | Basidiobolomycetes | *Basidiobolus ranarum* |
| ASV1614 | -2.663 | 7.24E-07 | Ascomycota | Dothideomycetes | *Pseudocercospora musae* |
| ASV1123 | -2.591 | 1.59E-05 | Ascomycota | Sordariomycetes | *Chaetomium globosum* |
| ASV4046 | -2.571 | 2.21E-06 | Ascomycota | Dothideomycetes | *Pseudocercospora musae* |
| ASV2899 | -2.438 | 8.80E-06 | Ascomycota | Saccharomycetes | *Pichia membranifaciens* |
| ASV617 | -2.309 | 1.56E-05 | Basidiobolomycota | Basidiobolomycetes | *Basidiobolus ranarum* |
| ASV1855 | -2.299 | 1.60E-05 | Ascomycota | Dothideomycetes | *Pseudocercospora musae* |

| | | | | | |
|---|---|---|---|---|---|
| ASV941 | -1.941 | 0.0008678114413 | Ascomycota | Sordariomycetes | *Niesslia tenuis* |
| ASV10350 | -1.876 | 0.0001613280111 | Ascomycota | Dothideomycetes | *Neodevriesia lagerstroemiae* |
| ASV10380 | -1.741 | 0.0004139051461 | Basidiomycota | Agaricomycetes | *Coprinellus radians* |
| ASV221 | -1.719 | 0.0008134754037 | Ascomycota | Sordariomycetes | *Chaetomium globosum* |
| ASV7033 | -1.508 | 0.0007735688578 | Basidiomycota | Agaricomycetes | *Bovista plumbea* |
| ASV942 | -1.507 | 0.0004459787692 | Ascomycota | Dothideomycetes | *Neodevriesia lagerstroemiae* |
| ASV2479 | 1.948 | 0.0001826917427 | Basidiomycota | Tremellomycetes | *Cryptococcus neoformans* |
| ASV54 | 2.359 | 1.88E-05 | Ascomycota | Eurotiomycetes | *Penicillium chrysogenum* |
| ASV2585 | 2.994 | 0.00013333795 | Basidiomycota | Microbotryomycetes | *Rhodotorula mucilaginosa* |
| ASV823 | 3.061 | 0.0002251994007 | Ascomycota | Dothideomycetes | *Alternaria citrimacularis* |
| ASV488 | 3.325 | 0.0001506284586 | Ascomycota | Sordariomycetes | *Chaetomium globosum* |
| ASV184 | 3.453 | 1.70E-05 | Basidiomycota | Tremellomycetes | *Naganishia vishniacii* |
| ASV371 | 3.482 | 0.0001993858026 | Ascomycota | Saccharomycetes | *Pichia membranifaciens* |
| ASV1576 | 3.590 | 1.26E-05 | Ascomycota | Dothideomycetes | *Alternaria tellustris* |
| ASV72 | 3.605 | 6.79E-05 | Ascomycota | Saccharomycetes | *Pichia membranifaciens* |
| ASV521 | 3.619 | 3.74E-05 | Ascomycota | Eurotiomycetes | *Exophiala heteromorpha* |
| ASV176 | 3.740 | 3.31E-06 | Basidiomycota | Microbotryomycetes | *Phenoliferia glacialis* |
| ASV279 | 3.819 | 4.56E-05 | Basidiomycota | Tremellomycetes | *Naganishia vishniacii* |
| ASV493 | 3.865 | 1.99E-05 | Basidiobolomycota | Basidiobolomycetes | *Basidiobolus ranarum* |
| ASV261 | 3.954 | 1.40E-06 | Basidiomycota | Microbotryomycetes | *Rhodotorula paludigena* |
| ASV756 | 4.147 | 5.31E-06 | Ascomycota | Sordariomycetes | *Claviceps purpurea* |

| | | | | | |
|---|---|---|---|---|---|
| ASV7356 | 4.331 | 2.37E-07 | Basidiomycota | Microbotryomycetes | *Rhodotorula mucilaginosa* |
| ASV422 | 4.825 | 8.15E-09 | Basidiomycota | Microbotryomycetes | *Rhodotorula mucilaginosa* |
| ASV717 | 4.910 | 5.94E-09 | Ascomycota | Dothideomycetes | *Alternaria tellustris* |
| ASV230 | 4.914 | 2.33E-08 | Ascomycota | Dothideomycetes | *Alternaria longipes* |
| ASV784 | 5.135 | 4.05E-09 | Ascomycota | Dothideomycetes | *Alternaria tellustris* |
| ASV688 | 5.160 | 3.36E-08 | Ascomycota | Dothideomycetes | *Alternaria tellustris* |
| ASV232 | 5.565 | 4.35E-11 | Basidiomycota | Microbotryomycetes | *Rhodotorula paludigena* |
| ASV249 | 5.576 | 7.81E-11 | Basidiomycota | Microbotryomycetes | *Rhodotorula mucilaginosa* |
| ASV32 | 5.595 | 9.55E-09 | Ascomycota | Dothideomycetes | *Alternaria tellustris* |
| ASV2065 | 5.772 | 2.30E-11 | Ascomycota | Dothideomycetes | *Alternaria tellustris* |
| ASV125 | 5.928 | 1.50E-11 | Basidiomycota | Microbotryomycetes | *Rhodotorula mucilaginosa* |
| ASV172 | 6.010 | 7.05E-09 | Basidiomycota | Agaricomycetes | *Rhizoctonia solani* |
| ASV103 | 6.371 | 1.03E-08 | Ascomycota | Dothideomycetes | *Alternaria tellustris* |
| ASV115 | 6.449 | 1.67E-10 | Ascomycota | Eurotiomycetes | *Exophiala heteromorpha* |
| ASV10944 | 6.657 | 8.37E-11 | Ascomycota | Eurotiomycetes | *Penicillium citrinum* |
| ASV35 | 6.664 | 5.14E-13 | Ascomycota | Dothideomycetes | *Alternaria tellustris* |
| ASV141 | 6.893 | 1.58E-10 | Basidiomycota | Agaricomycetes | *Rhizoctonia solani* |
| ASV4 | 7.461 | 5.20E-14 | Ascomycota | Saccharomycetes | *Pichia membranifaciens* |

**Table S3:** List of fungal isolates from the study and their taxonomy.

| Closest taxonomy, NCBI BLAST | Percent identity | Location collected | Substra te | First report? |
|---|---|---|---|---|
| *Rhodotorula kratochvilovae* | 99.64% | Anza Borrego Research Station, Colorado Desert | Moss crust | First report in Sonoran Desert |
| *Naganishia diffluens* | 99.82% | Anza Borrego Research Station, Colorado Desert | Moss crust | First report in Sonoran Desert |
| *Naganishia liquefaciens* | 99.65% | CIMA Volcanic Field, Mojave National Preserve | Moss crust | First report in Mojave Desert |
| *Naganishia liquefaciens* | 99.66% | CIMA Volcanic Field, Mojave National Preserve | Moss Crust | First report in Mojave Desert |
| *Naganishia liquefaciens* | 99.48% | CIMA Volcanic Field, Mojave National Preserve | Moss Crust | First report in Mojave Desert |
| *Naganishia albida* | 99.49% | CIMA Volcanic Field, Mojave National Preserve | Moss Crust | First report in Mojave Desert |
| *Naganishia albida* | 99.65% | CIMA Volcanic Field, Mojave National Preserve | Moss Crust | First report in Mojave Desert |
| *Kurtzmanomyces shapotouensis* | 99.64% | CIMA Volcanic Field, Mojave National Preserve | Lichen Crust | First report in Mojave Desert |
| *Kurtzmanomyces shapotouensis* | 99.64% | CIMA Volcanic Field, Mojave National Preserve | Moss crust | First report in Mojave Desert |
| *Coniochaeta luteorubra* | 91.94% | Granite Mountains, Mojave Desert | Moss Crust | First report in Mojave Desert |
| *Coniochaeta sp.* | 92.56% | Granite Mountains, Mojave Desert | Moss Crust | First report in Mojave |

| | | | | |
|---|---|---|---|---|
| | | | | Desert |
| *Coniochaeta luteorubra* | 91.84% | Granite Mountains, Mojave Desert | Moss Crust | First report in Mojave Desert |
| *Coniochaeta luteorubra* | 92.51% | Granite Mountains, Mojave Desert | Moss Crust | First report in Mojave Desert |
| *Coniochaeta canina* | 89.50% | Granite Mountains, Mojave Desert | Moss Crust | First report in Mojave Desert |
| *Cystobasidium pallidum* | 99.67% | Granite Mountains, Mojave Desert | Moss Crust | First report in Mojave Desert |
| *Coniochaeta sp.* | 96.00% | Granite Mountains, Mojave Desert | Moss Crust | First report in Mojave Desert |
| *Coniochaeta luteorubra* | 92.44% | Granite Mountains, Mojave Desert | Moss Crust | First report in Mojave Desert |
| *Coniochaeta luteorubra* | 92.27% | Granite Mountains, Mojave Desert | Moss Crust | First report in Mojave Desert |
| *Coniochaeta luteorubra* | 92.74% | Granite Mountains, Mojave Desert | Moss Crust | First report in Mojave Desert |
| *Coniochaeta sp.* | 92.76% | Granite Mountains, Mojave Desert | Moss Crust | First report in Mojave Desert |
| *Sordariomycetes sp.* | 79.79% | Granite Mountains, Mojave Desert | Moss Crust | Not applicable |
| *Coniochaeta sp.* | 92.92% | Granite Mountains, Mojave Desert | Moss Crust | First report in Mojave Desert |
| *Aureobasidium melanogenum* | 99.81% | Granite Mountains, Mojave Desert | Lichen Crust | Not a first report |
| *Coniochaeta sp.* | 92.26% | Granite Mountains, Mojave Desert | Moss Crust | First report in Mojave Desert |
| *Naganishia friedmannii* | 99.66% | Granite Mountains, Mojave Desert | Moss Crust | First report in Mojave Desert |
| *Exophiala nigra* | 98.37% | Granite Mountains, Mojave | Moss | First report in |

| | | | | |
|---|---|---|---|---|
| | | Desert | crust | Mojave Desert |
| *Exophiala crusticola* | 99.64% | Granite Mountains, Mojave Desert | Moss crust | Not a first report |
| *Aureobasidium melanogenum* | 99.81% | Granite Mountains, Mojave Desert | Lichen Crust | Not a first report |
| *Aureobasidium pullans* | 99.81% | Granite Mountains, Mojave Desert | Lichen Crust | Not a first report |
| *Rhodotorula sphaerocarpa* | 94.17% | Granite Mountains, Mojave Desert | Moss crust | First report in Mojave Desert |
| *Naganishia randhawae* | 99.83% | Oasis De Los Osos, Colorado Desert | Moss crust | First report in Sonoran Desert |
| *Naganishia liquefaciens* | 99.83% | Oasis De Los Osos, Colorado Desert | Moss crust | First report in Sonoran Desert |
| *Naganishia randhawae* | 100.00% | Oasis De Los Osos, Colorado Desert | Moss crust | First report in Sonoran Desert |
| *Naganishia globosa* | 98.44% | Oasis De Los Osos, Colorado Desert | Moss crust | First report in Sonoran Desert |
| *Aureobasidium pullulans* | 99.81% | Oasis De Los Osos, Colorado Desert | Moss crust | First report in Sonoran Desert |
| *Naganishia friedmannii* | 98.44% | Oasis De Los Osos, Colorado Desert | Moss crust | First report in Sonoran Desert |
| *Naganishia onofrii* | 100% | Oasis De Los Osos, Colorado Desert | Moss crust | First report in Sonoran Desert |
| *Aureobasidium pullulans* | 100.00% | Oasis De Los Osos, Colorado Desert | Moss crust | Not a first report |
| *Naganishia onofrii* | 100.00% | Oasis De Los Osos, Colorado Desert | Moss crust | First report in Sonoran Desert |
| *Naganishia globosa* | 100.00% | Oasis De Los Osos, Colorado Desert | Moss crust | First report in Sonoran Desert |
| *Naganishia onofrii* | 100.00% | Oasis De Los Osos, Colorado Desert | Moss crust | First report in Sonoran |

| | | | | |
|---|---|---|---|---|
| | | | | Desert |
| *Naganishia diffluens* | 99.82% | Oasis De Los Osos, Colorado Desert | Moss crust | First report in Sonoran Desert |
| *Rhodotorula kratochvilovae* | 99.82% | Oasis De Los Osos, Colorado Desert | Moss crust | First report in Sonoran Desert |
| *Rhodosporidiobolus poonsookiae* | 100% | Oasis De Los Osos, Colorado Desert | Moss crust | First report in Sonoran Desert |
| *Naganishia diffluens* | 99.64% | Oasis De Los Osos, Colorado Desert | Moss crust | First report in Sonoran Desert |
| *Coniochaeta sp.* | 97% | Oasis De Los Osos, Colorado Desert | Moss crust | Not a first report |
| *Coniochaeta boothii* | 100% | Oasis De Los Osos, Colorado Desert | Moss crust | First report in Sonoran Desert |
| *Naganishia diffluens* | 100% | Oasis De Los Osos, Colorado Desert | Moss crust | First report in Sonoran Desert |
| *Naganishia randhawae* | 98% | Oasis De Los Osos, Colorado Desert | Moss crust | First report in Sonoran Desert |
| *Naganishia onofrii* | 100.00% | Oasis De Los Osos, Colorado Desert | Moss crust | First report in Sonoran Desert |
| *Curvibasidium nothofagi* | 95.83% | Torrey Pines California State Natural Reserve Site 2 | Moss Crust | First report in Torrey Pines |
| *Filobasidium magnum* | 89.19% | Torrey Pines California State Natural Reserve Site 2 | Moss Crust | First report in Torrey Pines |
| *Coniochaeta polymorpha* | 99.41% | Torrey Pines California State Natural Reserve, Site 1 | Moss Crust | First report in Torrey Pines |
| *Filobasidium magnum* | 99.66% | Torrey Pines California State Natural Reserve, Site 1 | Moss Crust | First report in Torrey Pines |
| *Filobasidium magnum* | 99.19% | Torrey Pines California State Natural Reserve, Site 1 | Moss Crust | First report in Torrey Pines |
| *Filobasidium magnum* | 99.66% | Torrey Pines California State Natural Reserve, Site 1 | Moss Crust | First report in Torrey Pines |
| *Filobasidium magnum* | 99.49% | Torrey Pines California State Natural Reserve, Site 1 | Moss Crust | First report in Torrey Pines |

| | | | | |
|---|---|---|---|---|
| *Filobasidium magnum* | 99.83% | Torrey Pines California State Natural Reserve, Site 1 | Moss Crust | First report in Torrey Pines |
| *Filobasidium magnum* | 99.83% | Torrey Pines California State Natural Reserve, Site 1 | Moss Crust | First report in Torrey Pines |
| *Filobasidium magnum* | 99.66% | Torrey Pines California State Natural Reserve, Site 1 | Moss Crust | First report in Torrey Pines |
| *Filobasidium magnum* | 99.65% | Torrey Pines California State Natural Reserve, Site 1 | Moss Crust | First report in Torrey Pines |
| *Papiliotrema terrestris* | 100.00% | Torrey Pines California State Natural Reserve, Site 1 | Lichen Crust | First report in Torrey Pines |
| *Papiliotrema terrestris* | 100.00% | Torrey Pines California State Natural Reserve, Site 1 | Lichen Crust | First report at Torrey Pines |
| *Vishniacozyma melezitolytica* | 94.25% | Torrey Pines California State Natural Reserve, Site 1 | Lichen Crust | First report at Torrey Pines |
| *Myriangium duriaei* | 95.50% | Torrey Pines California State Natural Reserve, Site 1 | Lichen Crust | First report at Torrey Pines |
| *Solicoccozyma keelungensis* | 93.83% | Torrey Pines California State Natural Reserve, Site 1 | Lichen Crust | First report at Torrey Pines |
| *Solicoccozyma phenolica* | 100.00% | Torrey Pines California State Natural Reserve, Site 1 | Lichen Crust | First report at Torrey Pines |
| *Papiliotrema terrestris* | 99.79% | Torrey Pines California State Natural Reserve, Site 1 | Lichen Crust | First report at Torrey Pines |
| *Papiliotrema terrestris* | 99.79% | Torrey Pines California State Natural Reserve, Site 1 | Lichen Crust | First report at Torrey Pines |
| *Myriangium duriaei* | 95.50% | Torrey Pines California State Natural Reserve, Site 1 | Lichen Crust | First report at Torrey Pines |
| *Myriangium duriaei* | 95.50% | Torrey Pines California State Natural Reserve, Site 1 | Lichen Crust | First report at Torrey Pines |
| *Myriangium duriaei* | 95.50% | Torrey Pines California State Natural Reserve, Site 1 | Lichen Crust | First report at Torrey Pines |
| *Myriangium duriaei* | 95.50% | Torrey Pines California State Natural Reserve, Site 1 | Lichen Crust | First report at Torrey Pines |

**Table S4:** List of fungal species with differential abundance before and after surface sterilization. Note, fungi that could not be identified to the species level were filtered from this table.

| ASV | log2FoldChange | pvalue | Phylum | Class | Species |
|---|---|---|---|---|---|
| ASV688 | -5.389 | 4.28E-13 | Ascomycota | Dothideomycetes | *Alternaria tellustris* |
| ASV10944 | -4.895 | 2.19E-08 | Ascomycota | Eurotiomycetes | *Penicillium citrinum* |
| ASV230 | -4.709 | 6.60E-12 | Ascomycota | Dothideomycetes | *Alternaria longipes* |
| ASV521 | -4.093 | 1.34E-08 | Ascomycota | Eurotiomycetes | *Exophiala heteromorpha* |
| ASV35 | -3.862 | 1.25E-05 | Ascomycota | Dothideomycetes | *Alternaria tellustris* |
| ASV1576 | -3.782 | 1.20E-08 | Ascomycota | Dothideomycetes | *Alternaria tellustris* |
| ASV103 | -3.719 | 0.0001050823836 | Ascomycota | Dothideomycetes | *Alternaria tellustris* |
| ASV717 | -3.435 | 1.21E-05 | Ascomycota | Dothideomycetes | *Alternaria tellustris* |
| ASV784 | -3.417 | 7.38E-06 | Ascomycota | Dothideomycetes | *Alternaria tellustris* |
| ASV2065 | -3.113 | 4.64E-05 | Ascomycota | Dothideomycetes | *Alternaria tellustris* |
| ASV72 | -2.926 | 1.28E-05 | Ascomycota | Saccharomycetes | *Pichia membranifaciens* |
| ASV823 | -2.556 | 1.87E-05 | Ascomycota | Dothideomycetes | *Alternaria citrimacularis* |
| ASV1527 | -2.318 | 0.0001011794496 | Ascomycota | Dothideomycetes | *Alternaria citrimacularis* |
| ASV4692 | -2.246 | 4.56E-05 | Ascomycota | Dothideomycetes | *Alternaria tenuissima* |
| ASV3938 | -2.143 | 5.70E-05 | Ascomycota | Dothideomycetes | *Alternaria tenuissima* |
| ASV1614 | -2.116 | 0.0002018079025 | Ascomycota | Dothideomycetes | *Pseudocercospora musae* |
| ASV10055 | -2.078 | 0.0002100644268 | Basidiomycota | Tremellomycetes | *Naganishia vishniacii* |
| ASV2899 | -2.041 | 0.0003382024634 | Ascomycota | Saccharomycetes | *Pichia membranifaciens* |
| ASV5073 | 1.964 | 0.0001384119181 | Basidiomycota | Microbotryomycetes | *Rhodotorula mucilaginosa* |

| | | | | | |
|---|---|---|---|---|---|
| ASV630 | 2.090 | 0.0003701897834 | Ascomycota | Dothideomycetes | *Macrophomina phaseolina* |
| ASV3181 | 2.503 | 7.85E-05 | Ascomycota | Lecanoromycetes | *Protousnea dusenii* |
| ASV3686 | 2.570 | 4.64E-06 | Basidiomycota | Microbotryomycetes | *Rhodotorula mucilaginosa* |
| ASV617 | 2.654 | 0.0002066851129 | Basidiobolomycota | Basidiobolomycetes | *Basidiobolus ranarum* |
| ASV405 | 3.040 | 6.07E-06 | Ascomycota | Lecanoromycetes | *Protousnea dusenii* |
| ASV5982 | 3.101 | 4.30E-05 | Basidiomycota | Microbotryomycetes | *Sporobolomyces roseus* |
| ASV589 | 3.139 | 3.77E-05 | Ascomycota | Sordariomycetes | *Neocosmospora falciformis* |
| ASV423 | 3.470 | 1.87E-06 | Ascomycota | Lecanoromycetes | *Protousnea dusenii* |
| ASV349 | 3.511 | 1.78E-05 | Basidiobolomycota | Basidiobolomycetes | *Basidiobolus ranarum* |
| ASV493 | 3.570 | 4.58E-06 | Basidiobolomycota | Basidiobolomycetes | *Basidiobolus ranarum* |
| ASV862 | 3.611 | 6.63E-07 | Basidiomycota | Microbotryomycetes | *Rhodotorula paludigena* |
| ASV147 | 4.837 | 6.05E-09 | Ascomycota | Dothideomycetes | *Alternaria tenuissima* |
| ASV225 | 5.483 | 1.96E-12 | Basidiomycota | Microbotryomycetes | *Rhodotorula paludigena* |

**Table S5:** List of fungal species with differential abundance between soil and moss biocrust samples. Note, fungi that could not be identified to the species level were filtered from this table.

| Column 1 | log2FoldChange | pvalue | Class | Species |
|---|---|---|---|---|
| ASV17 | -9.812 | 4.00E-46 | Sordariomycetes | *Niesslia exosporioides* |
| ASV219 | -6.779 | 2.17E-25 | Sordariomycetes | *Niesslia tenuis* |
| ASV987 | -6.166 | 4.36E-25 | Sordariomycetes | *Trichoderma asperellum* |
| ASV406 | -5.909 | 2.86E-22 | Sordariomycetes | *Chaetomium subspirilliferum* |
| ASV118 | -5.815 | 3.81E-19 | Sordariomycetes | *Diatrypella favacea* |
| ASV468 | -5.345 | 9.92E-19 | Sordariomycetes | *Chaetomium subspirilliferum* |
| ASV132 | -5.150 | 5.78E-19 | Sordariomycetes | *Chaetomium globosum* |

| | | | | |
|---|---|---|---|---|
| ASV319 | -4.890 | 9.10E-17 | Sordariomycetes | *Chaetomium globosum* |
| ASV203 | -4.814 | 5.76E-16 | Sordariomycetes | *Diatrypella favacea* |
| ASV848 | -4.758 | 2.28E-17 | Sordariomycetes | *Niesslia tenuis* |
| ASV2890 | -4.288 | 2.40E-15 | Sordariomycetes | *Chaetomium globosum* |
| ASV890 | -4.251 | 1.87E-15 | Eurotiomycetes | *Penicillium canescens* |
| ASV351 | -4.245 | 6.99E-14 | Sordariomycetes | *Diatrypella favacea* |
| ASV942 | -4.196 | 8.30E-15 | Dothideomycetes | *Neodevriesia lagerstroemiae* |
| ASV450 | -4.056 | 2.81E-13 | Dothideomycetes | *Alternaria tenuissima* |
| ASV72 | -3.987 | 2.40E-08 | Saccharomycetes | *Pichia membranifaciens* |
| ASV331 | -3.910 | 3.41E-10 | Leotiomycetes | *Blumeria graminis* |
| ASV176 | -3.578 | 3.99E-08 | Microbotryomycetes | *Phenoliferia glacialis* |
| ASV13 | -3.557 | 1.33E-13 | Eurotiomycetes | *Penicillium chrysogenum* |
| ASV215 | -3.392 | 6.83E-08 | Dothideomycetes | *Alternaria longipes* |
| ASV1014 | -3.269 | 4.05E-09 | Dothideomycetes | *Alternaria tenuissima* |
| ASV299 | -3.066 | 8.67E-10 | Tremellomycetes | *Naganishia vishniacii* |
| ASV840 | -3.024 | 7.52E-11 | Sordariomycetes | *Chaetomium subspirilliferum* |
| ASV154 | -2.909 | 1.22E-08 | Sordariomycetes | *Chaetomium globosum* |
| ASV486 | -2.885 | 5.31E-09 | Sordariomycetes | *Fusarium verticillioides* |
| ASV1272 | -2.856 | 1.59E-09 | Sordariomycetes | *Diatrypella favacea* |
| ASV77 | -2.837 | 3.61E-06 | Dothideomycetes | *Alternaria tenuissima* |
| ASV9081 | -2.715 | 1.16E-08 | Sordariomycetes | *Chaetomium globosum* |
| ASV958 | -2.704 | 4.69E-09 | Sordariomycetes | *Diatrypella favacea* |
| ASV825 | -2.662 | 7.48E-08 | Sordariomycetes | *Chaetomium globosum* |
| ASV383 | -2.643 | 4.31E-08 | Sordariomycetes | *Chaetomium globosum* |
| ASV2462 | -2.639 | 0.0001561683362 | Sordariomycetes | *Diatrypella favacea* |
| ASV707 | -2.619 | 1.34E-07 | Eurotiomycetes | *Aspergillus ustus* |
| ASV757 | -2.588 | 5.76E-08 | Sordariomycetes | *Chaetomium globosum* |
| ASV771 | -2.569 | 2.87E-08 | Leotiomycetes | *Podosphaera leucotricha* |
| ASV777 | -2.566 | 1.52E-07 | Sordariomycetes | *Niesslia tenuis* |
| ASV230 | -2.456 | 0.0002974258269 | Dothideomycetes | *Alternaria longipes* |

| | | | | |
|---|---|---|---|---|
| ASV393 | -2.452 | 4.36E-05 | Microbotryomycetes | *Phenoliferia glacialis* |
| ASV806 | -2.447 | 2.90E-07 | Sordariomycetes | *Chaetomium globosum* |
| ASV512 | -2.336 | 9.72E-07 | Eurotiomycetes | *Aspergillus latus* |
| ASV903 | -2.308 | 2.55E-07 | Dothideomycetes | *Alternaria brassicae* |
| ASV816 | -2.304 | 2.91E-07 | Microbotryomycetes | *Phenoliferia glacialis* |
| ASV4624 | -2.282 | 6.60E-07 | Eurotiomycetes | *Aspergillus niger* |
| ASV1099 | -2.154 | 3.78E-06 | Eurotiomycetes | *Penicillium griseofulvum* |
| ASV604 | -2.111 | 0.0007289080437 | Dothideomycetes | *Stemphylium vesicarium* |
| ASV2070 | -2.051 | 2.86E-06 | Sordariomycetes | *Chaetomium globosum* |
| ASV190 | -2.049 | 2.42E-06 | Sordariomycetes | *Chaetomium globosum* |
| ASV787 | -2.031 | 1.26E-05 | Eurotiomycetes | *Aspergillus parasiticus* |
| ASV712 | -1.990 | 9.03E-05 | Saccharomycetes | *Pichia membranifaciens* |
| ASV248 | -1.965 | 7.94E-06 | Dothideomycetes | *Alternaria tenuissima* |
| ASV1211 | -1.903 | 1.73E-05 | Sordariomycetes | *Niesslia tenuis* |
| ASV392 | -1.872 | 2.28E-05 | Eurotiomycetes | *Aspergillus chevalieri* |
| ASV894 | -1.760 | 3.68E-05 | Agaricomycetes | *Efibula tuberculata* |
| ASV1197 | -1.738 | 5.58E-05 | Agaricomycetes | *Bovista plumbea* |
| ASV1378 | -1.732 | 3.40E-05 | Dothideomycetes | *Neodevriesia lagerstroemiae* |
| ASV1435 | -1.698 | 2.57E-05 | Eurotiomycetes | *Penicillium ochrochloron* |
| ASV3151 | -1.694 | 5.02E-05 | Eurotiomycetes | *Exophiala heteromorpha* |
| ASV1535 | -1.668 | 9.05E-05 | Dothideomycetes | *Alternaria longipes* |
| ASV2934 | -1.641 | 3.88E-05 | Tremellomycetes | *Naganishia vishniacii* |
| ASV4305 | -1.637 | 4.07E-05 | Eurotiomycetes | *Aspergillus flavus* |
| ASV2095 | -1.609 | 3.73E-05 | Lichinomycetes | *Heppia africana* |
| ASV1779 | -1.602 | 5.71E-05 | Eurotiomycetes | *Penicillium aurantiogriseum* |
| ASV560 | -1.598 | 0.0005682999082 | Dothideomycetes | *Pseudocercospora musae* |
| ASV1484 | -1.598 | 0.0001693670861 | Eurotiomycetes | *Aspergillus ustus* |
| ASV6184 | -1.595 | 4.49E-05 | Eurotiomycetes | *Penicillium chrysogenum* |
| ASV5434 | -1.579 | 5.93E-05 | Agaricomycetes | *Serendipita communis* |

| | | | | |
|---|---|---|---|---|
| ASV2281 | -1.578 | 0.000107368643 | Eurotiomycetes | *Penicillium oxalicum* |
| ASV1052 | -1.574 | 0.0003959121447 | Dothideomycetes | *Alternaria tellustris* |
| ASV2647 | -1.569 | 0.0001441458408 | Sordariomycetes | *Plectosphaerella cucumerina* |
| ASV2885 | -1.524 | 0.0001544175547 | Eurotiomycetes | *Aspergillus sydowii* |
| ASV2857 | -1.495 | 7.19E-05 | Eurotiomycetes | *Aspergillus ustus* |
| ASV5513 | -1.485 | 0.0001735852569 | Sordariomycetes | *Chaetomium subspirilliferum* |
| ASV579 | -1.483 | 0.0003561706109 | Sordariomycetes | *Chaetomium globosum* |
| ASV786 | -1.480 | 0.0005326146745 | Agaricomycetes | *Rhizoctonia solani* |
| ASV2362 | -1.462 | 0.000237437652 | Eurotiomycetes | *Penicillium lanosum* |
| ASV1102 | -1.427 | 0.0003294904808 | Agaricomycetes | *Bovista plumbea* |
| ASV490 | -1.416 | 0.0005482172139 | Dothideomycetes | *Alternaria longipes* |
| ASV2776 | -1.416 | 9.49E-05 | Sordariomycetes | *Chaetomium globosum* |
| ASV3182 | -1.405 | 0.0002980247654 | Dothideomycetes | *Alternaria citrimacularis* |
| ASV1127 | -1.372 | 0.000626324732 | Eurotiomycetes | *Exophiala heteromorpha* |
| ASV4759 | -1.358 | 0.0004071583815 | Sordariomycetes | *Stilbocrea gracilipes* |
| ASV1446 | -1.319 | 0.001073134773 | Agaricomycetes | *Bovista plumbea* |
| ASV3545 | -1.317 | 0.0005274815779 | Sordariomycetes | *Chaetomium subspirilliferum* |
| ASV2722 | -1.313 | 0.0004886157594 | Agaricomycetes | *Bovista plumbea* |
| ASV2666 | -1.304 | 0.0006676165586 | Sordariomycetes | *Paramyrothecium roridum* |
| ASV1301 | -1.281 | 0.0009989034545 | Agaricomycetes | *Bovista plumbea* |
| ASV2817 | -1.271 | 0.0008375418494 | Eurotiomycetes | *Aspergillus ustus* |

| | | | | |
|---|---|---|---|---|
| ASV2078 | -1.265 | 0.001434666788 | Dothideomycetes | *Alternaria citrimacularis* |
| ASV709 | -1.253 | 0.001507354714 | Sordariomycetes | *Niesslia tenuis* |
| ASV5460 | -1.184 | 0.0009039190957 | Eurotiomycetes | *Penicillium griseofulvum* |
| ASV2732 | -1.182 | 0.001171178905 | Eurotiomycetes | *Aspergillus latus* |
| ASV3550 | -1.173 | 0.001397800089 | Dothideomycetes | *Alternaria citrimacularis* |
| ASV7096 | -1.138 | 0.001422614316 | Lecanoromycetes | *Protousnea dusenii* |
| ASV2177 | -1.128 | 0.00072514678 | Sordariomycetes | *Niesslia exosporioides* |
| ASV9414 | 1.268 | 0.0002388396999 | Dothideomycetes | *Alternaria citrimacularis* |
| ASV1614 | 1.306 | 0.001081149014 | Dothideomycetes | *Pseudocercospora musae* |
| ASV3938 | 1.335 | 0.000446347615 | Dothideomycetes | *Alternaria tenuissima* |
| ASV941 | 1.597 | 0.0001081294939 | Sordariomycetes | *Niesslia tenuis* |
| ASV162 | 1.658 | 0.0007069108169 | Dothideomycetes | *Cercosporella tinosporae* |
| ASV3805 | 1.696 | 7.80E-05 | Sordariomycetes | *Neocosmospora falciformis* |
| ASV1326 | 1.704 | 8.71E-05 | Dothideomycetes | *Neodevriesia lagerstroemiae* |
| ASV489 | 1.806 | 8.58E-05 | Glomeromycetes | *Glomus macrocarpum* |
| ASV630 | 1.830 | 6.16E-05 | Dothideomycetes | *Macrophomina phaseolina* |
| ASV405 | 1.845 | 7.44E-05 | Lecanoromycetes | *Protousnea dusenii* |
| ASV10149 | 2.171 | 1.57E-06 | Tremellomycetes | *Vishniacozyma victoriae* |
| ASV9 | 2.173 | 8.66E-05 | Sordariomycetes | *Neocosmospora falciformis* |
| ASV310 | 2.204 | 3.59E-06 | Agaricomycetes | *Hericium erinaceus* |
| ASV2479 | 2.209 | 4.17E-08 | Tremellomycetes | *Cryptococcus neoformans* |
| ASV7356 | 2.492 | 3.85E-06 | Microbotryomycetes | *Rhodotorula mucilaginosa* |
| ASV589 | 2.529 | 1.00E-06 | Sordariomycetes | *Neocosmospora falciformis* |
| ASV756 | 2.695 | 5.51E-08 | Sordariomycetes | *Claviceps purpurea* |

| | | | | |
|---|---|---|---|---|
| ASV488 | 3.543 | 8.41E-11 | Sordariomycetes | *Chaetomium globosum* |
| ASV349 | 3.598 | 1.82E-10 | Basidiobolomycetes | *Basidiobolus ranarum* |
| ASV1094 4 | 5.037 | 6.40E-16 | Eurotiomycetes | *Penicillium citrinum* |

Table S6: GenBank accessions for each isolate in the study.

| Accession | Sequence ID |
|---|---|
| PV082275 | JDC1 |
| PV082276 | JDC2 |
| PV082277 | JDC3 |
| PV082278 | JDC4 |
| PV082279 | JDC5 |
| PV082280 | JDC6 |
| PV082281 | JDC7 |
| PV082282 | JDC8 |
| PV082283 | JDC9 |
| PV082284 | JDC10 |
| PV082285 | JDC11 |
| PV082286 | JDC13 |
| PV082287 | JDC15 |
| PV082288 | JDC16 |
| PV082289 | JDC17 |
| PV082290 | JDC18 |
| PV082291 | JDC19 |
| PV082292 | JDC20 |

| | |
|---|---|
| PV082293 | JDC22 |
| PV082294 | JDC23 |
| PV082295 | JDC24 |
| PV082296 | JDC25 |
| PV082297 | JDC26 |
| PV082298 | JDC27 |
| PV082299 | JDC28 |
| PV082300 | JDC30 |
| PV082301 | JDC31 |
| PV082302 | JDC32 |
| PV082303 | JDC33 |
| PV082304 | KHK1 |
| PV082305 | KHK9 |
| PV082306 | KHK23 |
| PV082307 | KHK29 |
| PV082308 | KHK45 |
| PV082309 | KHK47 |
| PV082310 | KHK61 |
| PV082311 | KHK62 |
| PV082312 | KHK63 |
| PV082313 | KHK64 |
| PV082314 | KHK65 |
| PV082315 | KHK68 |

| | |
|---|---|
| PV082316 | KHK70 |
| PV082317 | KHK71 |
| PV082318 | KHK72 |
| PV082319 | KHK73 |
| PV082320 | KHK74 |
| PV082321 | KHK75 |
| PV082322 | KHK76 |
| PV082323 | KHK77 |
| PV082324 | KHK86 |
| PV082325 | KHK87 |
| PV082326 | KHK88 |
| PV082327 | VB1 |
| PV082328 | VB2 |
| PV082329 | VB5 |
| PV082330 | VB6 |
| PV082331 | VB9 |
| PV082332 | VB12 |
| PV082333 | VB13 |
| PV082334 | VB14 |
| PV082335 | VB15 |
| PV082336 | VB16 |
| PV082337 | VB17 |
| PV082338 | VB19 |

| | |
|---|---|
| PV082339 | VB21 |
| PV082340 | KHK59 |
| PV082341 | VB_infinity |
| PV082342 | KHK91 |
| PV082343 | KHK92 |
| PV082344 | KHK93 |
| PV082345 | KHK39 |
| PV082346 | AB-10-1 |
| PV082347 | VB18 |
